# Hippocampo-septal inhibitory feedback controls the medial septal circuit as a function of brain state, pyramidal cell activity and movement speed

**DOI:** 10.64898/2026.09.03.749177

**Authors:** Andor Domonkos, Dániel Schlingloff, Bálint Király, Litsa Ledri, Márta Jelitai, Gergely Turi, Mária Rita Karlocai, Edit Papp, Panna Hegedüs, Gábor Nyíri, Attila I. Gulyás, Tamás F. Freund, Karl Deisseroth, Attila Losonczy, Balázs Hangya, Viktor Varga

## Abstract

Encoding novel experiences and retrieving stored information rely on the hippocampal theta rhythm in several mammalian species. Coordination of putative pacemakers in the medial septum (MS) by the GABAergic hippocampo-septal (HS) feedback is hypothesized to be essential for theta genesis. In order to reveal how the HS-feedback controls the MS circuit, and through it hippocampal oscillations in general, we set out to uncover the behavior-related activity of HS neurons as well as how different components of the MS network are regulated by the HS-feedback. Utilizing computational modelling and in vivo optogenetic manipulation of the HS pathway, we show that the HS feedback is not required for theta generation, which is primarily governed by intra-MS connections. HS blockade did not alter the phase coupling of MS neurons to hippocampal theta rhythm nor the synchrony between rhythmic MS neurons but augmented ongoing theta and coupled mid-gamma oscillations in the hippocampus. By contrast, correlated activity between non-rhythmic MS neurons was sensitive to HS blockade. Moreover, the HS feedback scaled with hippocampal activity and running speed. In turn, modulation of MS neurons by running and ripples was diminished by the transient suppression of HS terminals. These findings suggest that the HS feedback exerts inhibitory control of the MS depending upon hippocampal output and movement speed.

## INTRODUCTION

Theta oscillations organize information flow in episodic memory circuits, dominating hippocampal activity during exploration and REM sleep and coordinating neuronal ensembles during encoding^1–3^. Precisely timed recruitment of neurons in the hippocampus is essential for generating temporally structured ensemble activity and for stabilizing hippocampal representations that support learning and memory^4–6^. The medial septum (MS) is a central theta pacemaker^7–9^, providing GABAergic, cholinergic and glutamatergic input^10–13^ that entrains hippocampal network activity supporting spatial, contextual and episodic memories^14–18^.

MS neurons can intrinsically generate theta-frequency bursts^19^, and septal inactivation disrupts hippocampal assemblies and impairs memory^20–23^. However, the MS and the hippocampus form a reciprocal loop^24–27^: hippocampo-septal (HS) projections can strongly influence septal theta generation and may rival or exceed the septo-hippocampal drive^28–30^. HS-projecting neurons constitute a small, heterogeneous and poorly characterized GABAergic population in the hippocampus. In dorsal CA1, most HS-projecting cells reside in stratum oriens, co-express somatostatin (SST) and GABA, receive unusually dense excitatory input from principal cells and target local interneurons, pyramidal cells, the MS and the supramammillary nucleus, among other regions^26,31–33^. These neurons have been reported to fire irregularly while remaining phase-locked to theta^31^ or to be selectively engaged during ripples^34,35^. Their extensive excitatory drive and prominent septal output suggest that they could critically regulate hippocampal rhythmogenesis and oscillatory coupling. However, due to the lack of data concerning both the *in vivo* behavior-dependent activity of HS neurons and the effect of their output on the MS rhythm generator, the role of this prominent inhibitory feedback in the formation of memory-related hippocampal patterns is still elusive. Moreover, this knowledge gap precluded the development of a working theory explaining rhythmogenesis in the septo-hippocampo-septal loop.

To unravel the function of HS feedback, we devised a multipronged strategy. First, to explore how HS neurons are recruited during behavior, we performed calcium imaging of retrogradely labeled HS cells in awake animals. Then, we determined how pyramidal cells’ activity affects the output of HS neurons using acute slice electrophysiology. Further, we tested how the HS feedback influences theta generation by computational modeling and selective optogenetic manipulation of the HS pathway in anesthetized and freely behaving mice. We uncovered that HS neurons projecting to the MS encode running speed and dynamically convert graded hippocampal excitation into state-dependent inhibition of MS neurons, thereby modulating - but not abolishing - septal theta rhythmic output. These results identify the HS pathway as a main feedback controller that tunes septo-hippocampal coordination according to ongoing hippocampal network demands rather than acting as a primary theta generator, revealing a novel circuit motif that flexibly reshapes oscillatory dynamics to meet current computational needs.

## RESULTS

### Activity of HS neurons is correlated with running speed

To monitor HS activity *in vivo,* we injected G-protein deleted rabies-GCaMP6f constructs^36^ (see Methods) into the MS, enabling retrograde labeling and imaging of HS neurons in head-restrained mice during treadmill running (Fig. 1a,b). HS neurons were primarily located in stratum oriens of dorsal CA1 and expressed SST^31^ (Fig. 1b). Occasional labeling of pyramidal cells was observed (Fig. 1b), likely due to limited viral spread into adjacent lateral septum^37^ or transduction of recently described glutamatergic hippocampal neurons projecting to the MS^38^. The analysis was restricted therefore to neurons located in stratum oriens.

**Figure 1.**
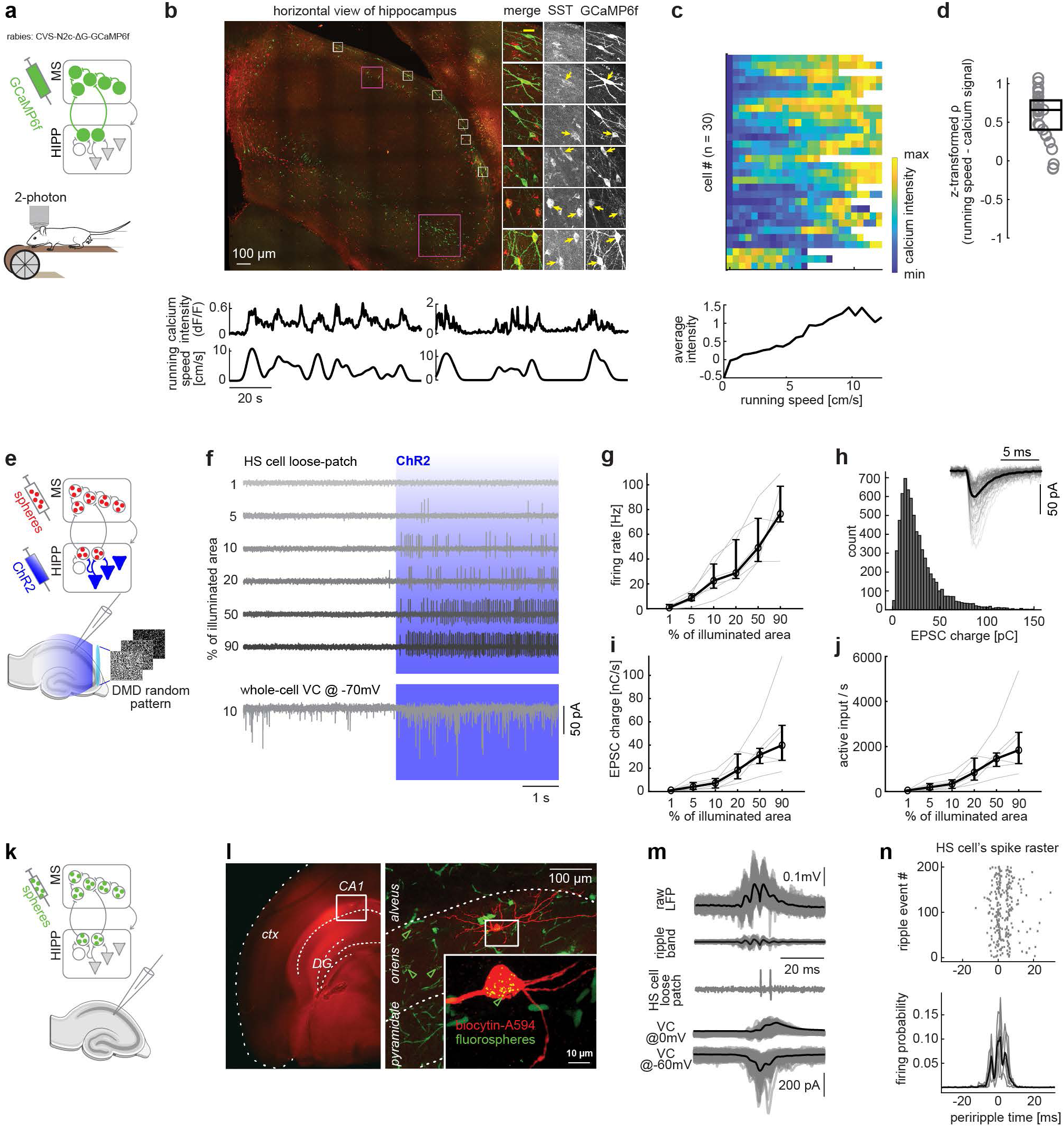
Recruitment of HS neurons. **a,** Schematic of the 2-photon imaging experiments. HS neurons were retrogradely transduced to express GCaMP6f using CVS-N2c^ΔG^ (n = 2 mice) or EnvA-pseudotyped CVS-N2c^ΔG^ (n = 2 mice) rabies vectors injected into the MS of wild type or Pvalb-Cre mice; for pseudotyped rabies, rAAV1/2(CAG- TVA-mCherry)Cre and rAAV1/2[EF1a-histoneGFP- 2A-rabiesG(N2c)]Cre helper viruses were co-injected into the MS. Imaging was performed in the hippocampus (HIPP) of head-restrained mice running on a treadmill. **b,** Top, horizontal view of the imaged hippocampal dorsal CA1 area; white boxes indicate example HS cells in stratum oriens (right, enlarged; immunolabeling for somatostatin (SST) and GCaMP6f); yellow arrows mark SST-positive and GCaMP6f-cells. Note that some principal cells (magenta boxes) were also labelled by GCaMP6f, these were not included in this study. Bottom, motion-corrected calcium traces of two demonstrative HS cells and simultaneously recorded running speed, illustrating up to 200% fluorescence increases during locomotion. **c,** Mean calcium signal intensity of individual HS cells as a function of running speed. Cells are ordered by ascending Spearman’s ρ (speed vs. signal intensity). Note that the calcium intensity strongly followed the speed increment in the majority of neurons, except n = 3 cells, in which the signal dropped by speed. **d,** Fisher ρ-z transformed Spearman’s ρ between running speed and calcium signal for individual HS cells. Box plot shows median and interquartile range; population-level correlation estimate was 0.5264 (p = 3.1817×10^-6^). Circles are ordered (left to right) by ascending p-value. **e,** Schematic of the in vitro experimental configuration. To examine HS cell responses to increasing pyramidal cell input, AAV1.CaMKIIα.hChR2(H134R)-mCherry.WPRE.hGH was injected into the hippocampus, and fluorescent microspheres into MS to identify HS cells. Labeled HS cells were recorded in acute slices, while excitatory input was recruited incrementally using a DMD-based patterned illumination with increasing pixel saturation of randomized light spots. **f,** Example of a HS cell recorded in loose patch mode showing spiking responses to increasing illumination levels (top to bottom, numbers indicate the pixel saturation over ∼180µm x 180µm illuminated area); the same neuron was subsequently recorded in whole-cell voltage clamp at -70 mV to measure EPSCs (bottom trace, 10 % illumination). **g,** Evoked HS cell firing rates plotted against illumination level (black, mean; grey traces, individual cells, n=6). **h,** Distribution of unitary EPSC charge; inset, individual EPSCs, the average shown in black. (Median: 21.66 (Q1:13.35, Q3:35.35) in pC, ∼7000 EPSCs from n=6 cells). **i-j,** Cumulative EPSC charge transfer (i) across illumination levels, and estimated number of active excitatory inputs (j), obtained by dividing cumulative charge transfer by the median unitary EPSC charge; HS cells approximately linearly transform excitatory synaptic input into action potential output. **k,** Retrograde labeling of hippocampal neurons projecting to the MS by fluorosphere injection, followed by recordings in acute slices. **l,** Confocal image of a recorded HS cell in the stratum oriens of ventral CA1 (biocytin-fill revealed with streptavidin-Alexa Fluor 594); fluorospheres injected into the MS accumulate in the HS somata (green arrowhead). **m,** Hippocampal LFP (raw and ripple band filtered traces aligned to ripples) and simultaneously recorded HS cell in loose-patch configuration (middle) and voltage clamp to reveal average IPSCs and EPSCs (bottom) in acute slice. **n,** Top, spike raster of an example HS cell aligned to sharp wave ripple events. Bottom, periripple time histogram (aligned to ripple peaks) of n = 5 loose patch recorded HS cells; black trace indicates the population mean.

The calcium signals of HS neurons closely followed running speed (Fig. 1b), showing robust fluorescence increases with increasing speed and revealing a broad dynamic range of HS activity modulation (Fig. 1c,d). A small subset (3 out of 30 neurons) showed inverse correlations with speed, suggesting a heterogeneity in how different HS cells report locomotor – or network – state (Fig. 1c,d).

### Recruitment of HS neurons by pyramidal cells

The broad modulation of HS activity observed *in vivo* suggested that these neurons, which collect extensive excitatory input from hippocampal pyramidal cells^32,33^, may track population-level excitatory activity. Hence, we next examined how HS firing scales with pyramidal cell drive in acute slices, where excitatory drive can be precisely controlled. We recorded from retrobead-labeled HS neurons while incrementally increasing the size of the optogenetically activated population of ChR2-expressing CA1 pyramidal cells via patterned illumination (Fig. 1e).

Increasing pyramidal cell output led to a graded increase in HS firing rates (Fig. 1f,g). Analysis of unitary EPSC charge measured in HS cells (Fig. 1h) revealed that stronger illumination recruited proportionally more excitatory inputs, resulting in an approximately linear increase in cumulative synaptic charge and output firing (Fig. 1i,j). These findings indicate that HS neurons transform hippocampal excitatory population activity into their spiking output in a quasi-linear manner.

### Cholinergic modulation of HS-neurons

Given that acetylcholine is a key movement-regulated modulator^39^ and profoundly reshapes hippocampal network dynamics^40–42^, we tested how increased cholinergic tone affects HS neuron activity. Bath application of the muscarinic agonist carbachol robustly increased HS firing in acute slices (Extended Data Fig. 1a-c). However, in the presence of ionotropic glutamate receptor blockers (NBQX, AP-5), carbachol produced only modest changes in HS firing and did not appreciably alter the resting membrane potential (Extended Data Fig. 1d,e). These observations suggest that cholinergic activation of the hippocampal network influences HS neurons largely indirectly through modulation of pyramidal cells, which in turn can reliably drive HS firing.

### Ripple-related recruitment of HS neurons

Sharp-wave ripples are generated by prominent, highly synchronized discharges of CA1 pyramidal cells and certain interneurons^43^, which possibly also recruit the long-range projecting HS neurons^34,35^. Consistent with this, retrobead-labeled HS neurons (Fig. 1k,l) showed coordinated spiking and synaptic currents aligned to spontaneous ripple-like transients in acute slices (Fig. 1m,n). HS activity increased several-fold during ripples compared to inter-ripple periods (Fig. 1n).

Together, the broad modulation range of HS neurons during locomotion, their graded recruitment by controlled optogenetic activation of pyramidal cells, their indirect cholinergic modulation and their ripple-coupled firing demonstrate that HS neurons are able to closely track hippocampal activity across behavioral states, suggesting a role that extends beyond the classical view of HS cells as components of the septo-hippocampo-septal theta-generating loop^24^.

### *In silico* model for HS impact on the MS

To explore how the dynamically modulated HS input can influence the MS circuitry and theta generation, we expanded a recently implemented *in silico* model of the MS network^19^ with incorporating the HS feedback. In this model, GABAergic neurons endowed with characteristic intrinsic conductances (Fig. 2a) were interconnected via fast GABAA synapses with a preset probability, and HS feedback was represented as an additional GABAA input targeting MS neurons (Fig. 2b). Tonic excitation synchronized MS neurons and generated theta rhythmic output (Fig. 2b,c), while MS neurons segregated into two populations: followers with theta-rhythmic firing only during induced theta state, and pacemakers with constant theta-rhythmicity^19^.

**Figure 2.**
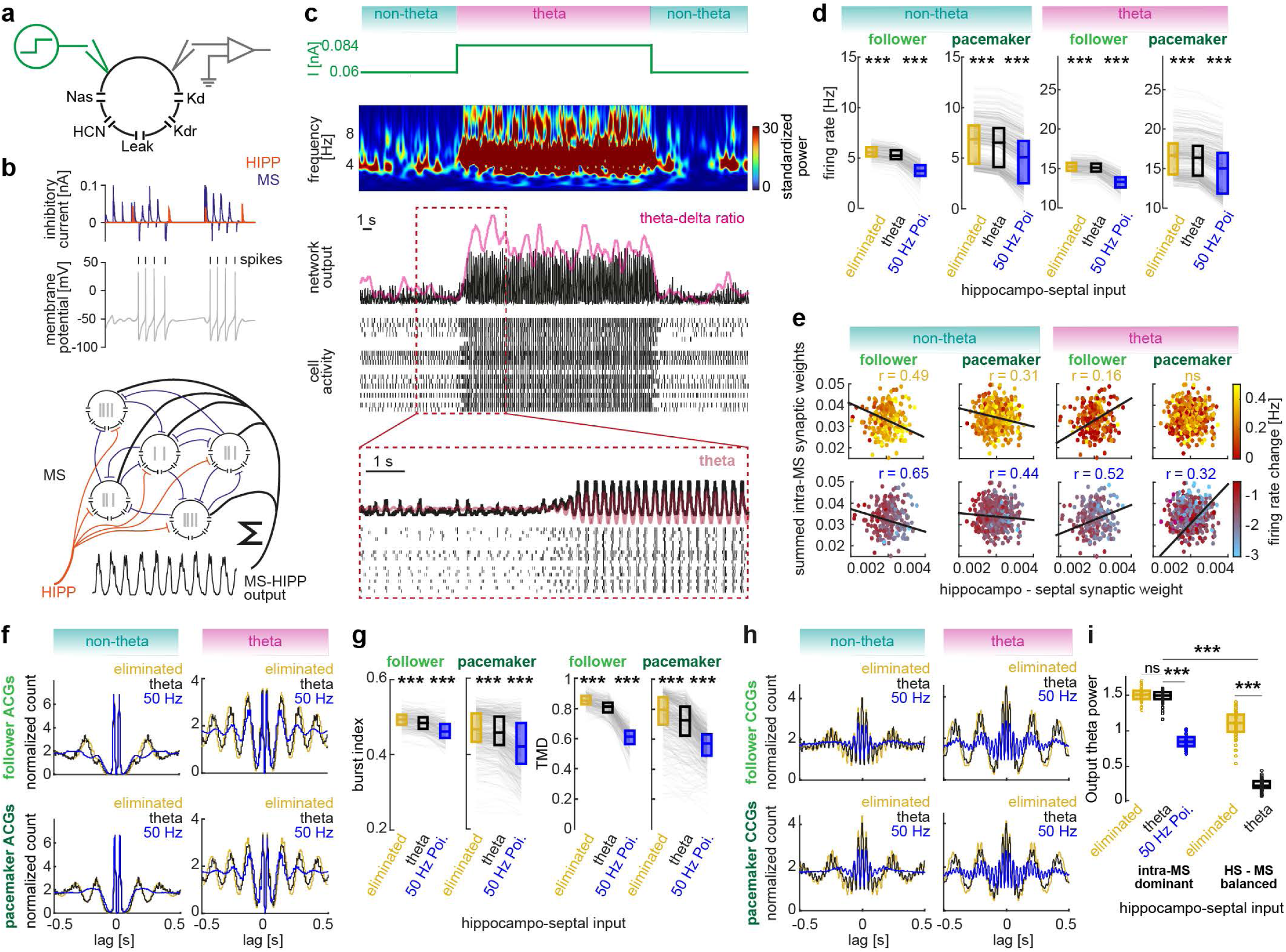
*In silico* model for HS impact on the MS. **a,** Schematic of the single-compartment inhibitory neuron model, including transient sodium (Nas), slowly inactivating D-type potassium (Kd), delayed-rectifier potassium (Kdr), HCN and passive leak channels. Tonic excitatory currents are delivered to the neuron and the membrane potential is recorded. **b,** Schematic of the septal network model. Twenty septal neurons receive GABA-A-mediated inhibitory synaptic input from the hippocampus and are also interconnected with a 60% connection rate (7 ms mean synaptic delay, 2 ms mean synaptic decay and 3 nS mean synaptic strength, each with 10% variance). The membrane potential of each cell is recorded to detect individual spike trains (top). Network output is computed as the sum of all spike trains convolved with a 50-ms Gaussian kernel (bottom). **c,** Simulation of network dynamics. The network is initially driven by a baseline current of 60 pA (with 10% variance) which is then increased to 84 pA to induce a highly synchronous theta state. This synchrony diminishes when the current is reset to baseline, reverting the network to a desynchronized (non-theta) state. Top, wavelet spectrogram of the network output. Middle, network output (black, raw; magenta, theta-delta power ratio) and simulated spike raster. Bottom, magnified segment around the increase in stimulation, showing the emergence of network synchrony (black, raw; red, theta-filtered signal; bottom, spike raster). The full simulation was repeated 60 times under three hippocampal input conditions: baseline theta-rhythmic (shown in this panel), fully suppressed, and 50 Hz Poisson-like activation. **d,** Elimination and activation of the HS input slightly but consistently increased and decreased, respectively, the firing rates of both pacemaker and follower model neurons across theta and non-theta states (***p < 0.001, paired Wilcoxon signed-rank test). Pacemaker and follower model neurons were defined based on their theta and non-theta state rhythmicity, analogous to the classification of real neurons. Box plots show median and interquartile range. **e,** HS input elimination (top) and activation (bottom) evoked a firing rate change of single model neurons as a function of the hippocampal and the summed intra-MS inhibitory synaptic weights. Trend lines indicate the gradient direction derived from a linear fit; r values reflect model fit quality, expressed as the correlation between actual and predicted firing rate changes based on this linear combination of synaptic weights. **f,** Average autocorrelograms of pacemaker and follower model neurons during non-theta and theta states, under suppressed, baseline and activated HS input conditions. **g,** Burst index (left) and theta modulation (right) increased with elimination and decreased with activation of the HS input (***p < 0.001, paired Wilcoxon signed-rank test). Box plots show median and interquartile range. **h,** Average cross-correlograms between pacemaker (top) and follower (bottom) model neurons during non-theta (left) and theta (right) states, under eliminated, baseline and activated HS input conditions. **i,** Left, theta power of the network output was not significantly altered by HS elimination, but significantly decreased when the input was activated (***p < 0.001, Mann-Whitney test). Right, theta power of the network output is significantly smaller in a network with balanced summed intra-MS and HS input strengths than in the intra-MS-connection-dominated network of panel b; elimination of the HS feedback in this balanced network robustly elevated the theta power of the network output (***p < 0.001, Mann-Whitney test).

Theta and non-theta states were simulated under three HS feedback regimes: theta-rhythmic (baseline) input contrasted with faster non-rhythmic (50 Hz Poisson-patterned) input or with the lack of HS input (feedback eliminated). Feedback elimination produced a modest but highly significant firing rate increase of both followers and pacemakers, in contrast to the large activity drop caused by the 50 Hz Poisson input (Fig. 2d).

We next examined how this effect depended on connectivity, mapping the HS-evoked firing rate changes of single neurons against both the HS synaptic weight and the summed intra-MS synaptic weight. During non-theta state, as expected, the HS synaptic weight was a strong determinant of the magnitude of the feedback effect, and weaker intra-MS connectivity was associated with a larger impact. During theta state, however, connection weights were weaker determinants (note the lack of significant relationship between HS and intra-MS synaptic weights in pacemaker neurons), and neurons with comparably strong intra-MS and HS connections were the most affected (Fig. 2e).

Overall, rhythmicity and synchrony among MS neurons during induced theta state were relatively preserved under both elimination and activation of the HS feedback (Fig. 2f-h), with a slight increase in burst index and theta modulation upon HS feedback elimination and a decrease upon activation (Fig. 2g), indicating that the intrinsic MS circuit is sufficient to generate theta without HS feedback. Accordingly, when intrinsic connections dominated the MS circuit, HS suppression had minimal impact on theta power, and even in a configuration with balanced HS and intra-MS connectivity – characterized by reduced baseline theta output –, HS suppression increased theta. Conversely, injecting high-frequency Poisson-patterned inhibition into the MS via HS feedback significantly reduced theta output (Fig. 2i).

Thus, computer simulations support a division of labor between intra-MS connections and HS feedback: intra-MS circuitry primarily sets theta rhythmic output through inhibitory synchronization mechanisms, while HS feedback acts as an inhibitory control that scales with hippocampal output.

### Optogenetic testing of HS impact on the MS

To examine the impact of the HS pathway on MS neurons *in vivo*, we combined optogenetic manipulations of HS terminals with juxtacellular or silicon probe recordings of activity in the MS (Fig. 3a, Fig. 4a). Using Sst-Cre mice for transduction of HS neurons to express the optogenetic actuators, we observed minimal off-target labeling beyond hippocampal SST neurons, and these events did not measurably influence the overall effects of our manipulations (Fig. 3b, Extended Data Fig. 2). To avoid modulation of hippocampal neurons and connections that do not target the MS^44^, illumination was restricted to the MS or the fimbria, where HS axons pass toward the septum. We first characterized the effects of optogenetic activation (Fig. 3) and inhibition (Fig. 4) under anesthesia with juxtacellular single-cell recordings (Fig. 3c,d), followed by chronic silicon probe recordings of multiple MS neurons in freely behaving mice (Fig. 3e).

**Figure 3.**
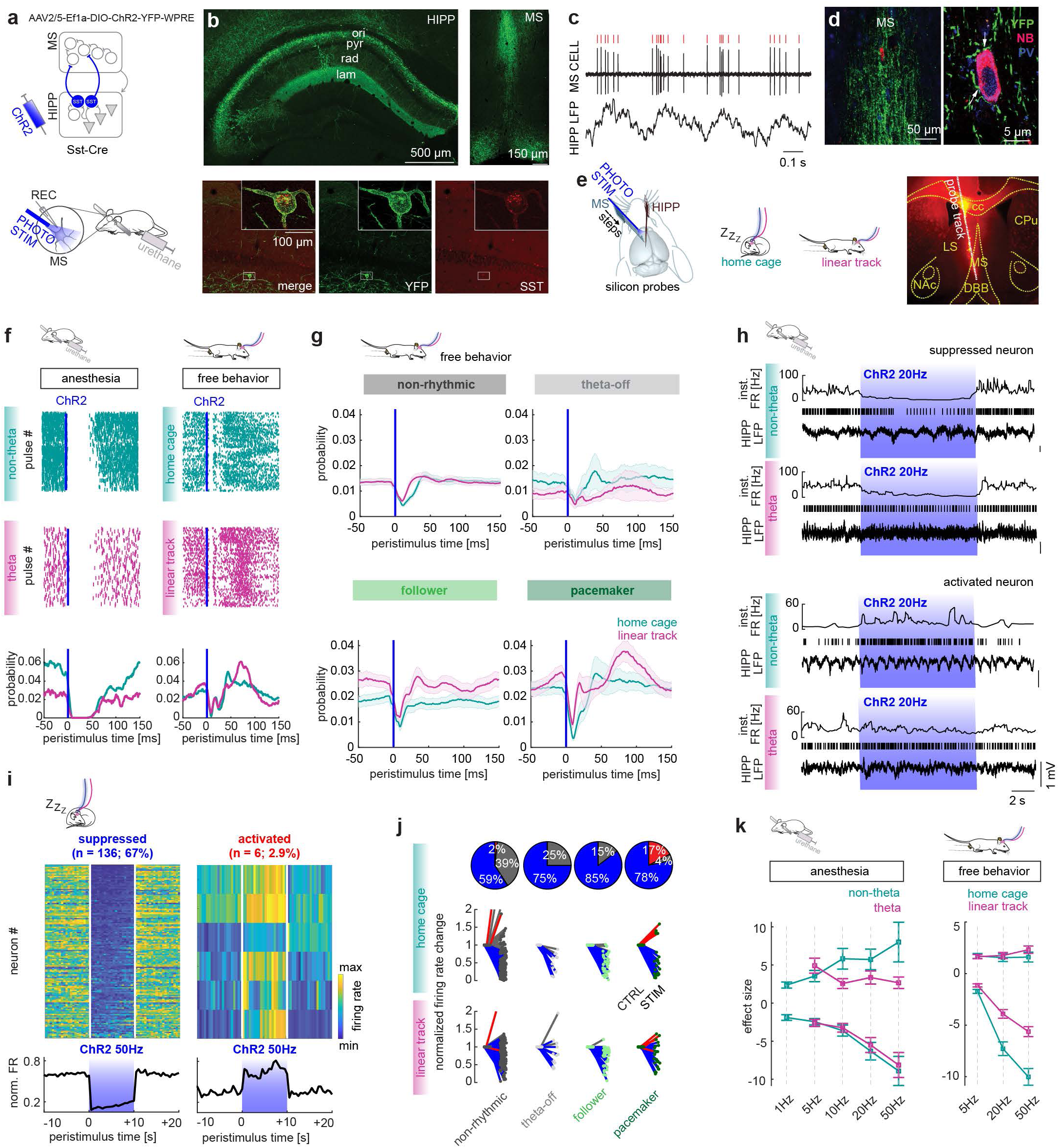
Optogenetic activation of the HS feedback. **a,** Experimental design for optical activation of the HS pathway. A Cre-dependent virus was injected into the hippocampus of Sst-Cre mice. Bottom, design of juxtacellular recording with optogenetic manipulation of HS pathway of MS neurons under urethane anesthesia. **b,** Top left, YFP expression in the hippocampus following viral injection. SST-positive cells were mainly located in the stratum oriens, extending axonal arbor into the stratum lacunosum-moleculare (indicating that O-LM cells are among the transduced neurons). Top right, YFP-labeled fibers were also observed in the MS, confirming that hippocampal SST neurons projecting to the septum were transduced. Bottom, magnified view of a YFP-expressing SST(+) neuron. **c,** Representative example of juxtacellular recording from an MS neuron (top, red ticks mark spike times) simultaneously with hippocampal LFP (bottom). **d,** Left, localization of the recorded MS neuron surrounded by YFP-positive hippocampal fibers. Right, magnified view. NB: neurobiotin, PV parvalbumin. **e,** Chronic recording design: silicon probes were implanted into the MS and hippocampus. Recordings were performed in home cage (quiet periods) and on a linear track (locomotion). Right, histological reconstruction of the probe track within the MS. **f,** Top, Spike raster plots during hippocampal non-theta or home cage (top) and theta or linear track (middle) states for an example pacemaker cell recorded juxtacellularly in anesthesia (left) or chronically by silicon probe in free behavior (right), aligned to 5-ms ChR2 pulses repeated at 5 Hz. Bottom, peristimulus time histograms (PSTHs, 1-ms bins) showing firing probability around the optical stimulation. **g,** Group average PSTHs (±SEM) in free behavior for the rhythmicity cell groups. **h,** Effect of 20 Hz ChR2 stimulation trains on MS firing under anesthesia. The upper example shows suppression, the lower example shows activation. Illumination epochs are indicated by shaded boxes. **i,** Effect of 50 Hz ChR2 stimulation trains on MS firing in free behavior, in home cage. The normalized firing rates of significantly suppressed (left) or activated (right) neurons are shown. Bottom, population average. **j**, Top, Proportion of suppressed (blue) and activated (red) neurons across rhythmicity groups in home cage (percentages rounded). Bottom, firing rate change relative to control, in home cage and in linear track. **k,** Effect size (firing rate during stimulation minus control rate) for increasing ChR2 stimulation frequencies (left, urethane anesthesia; right, free behavior). Positive and negative effect sizes are shown separately.

**Figure 4.**
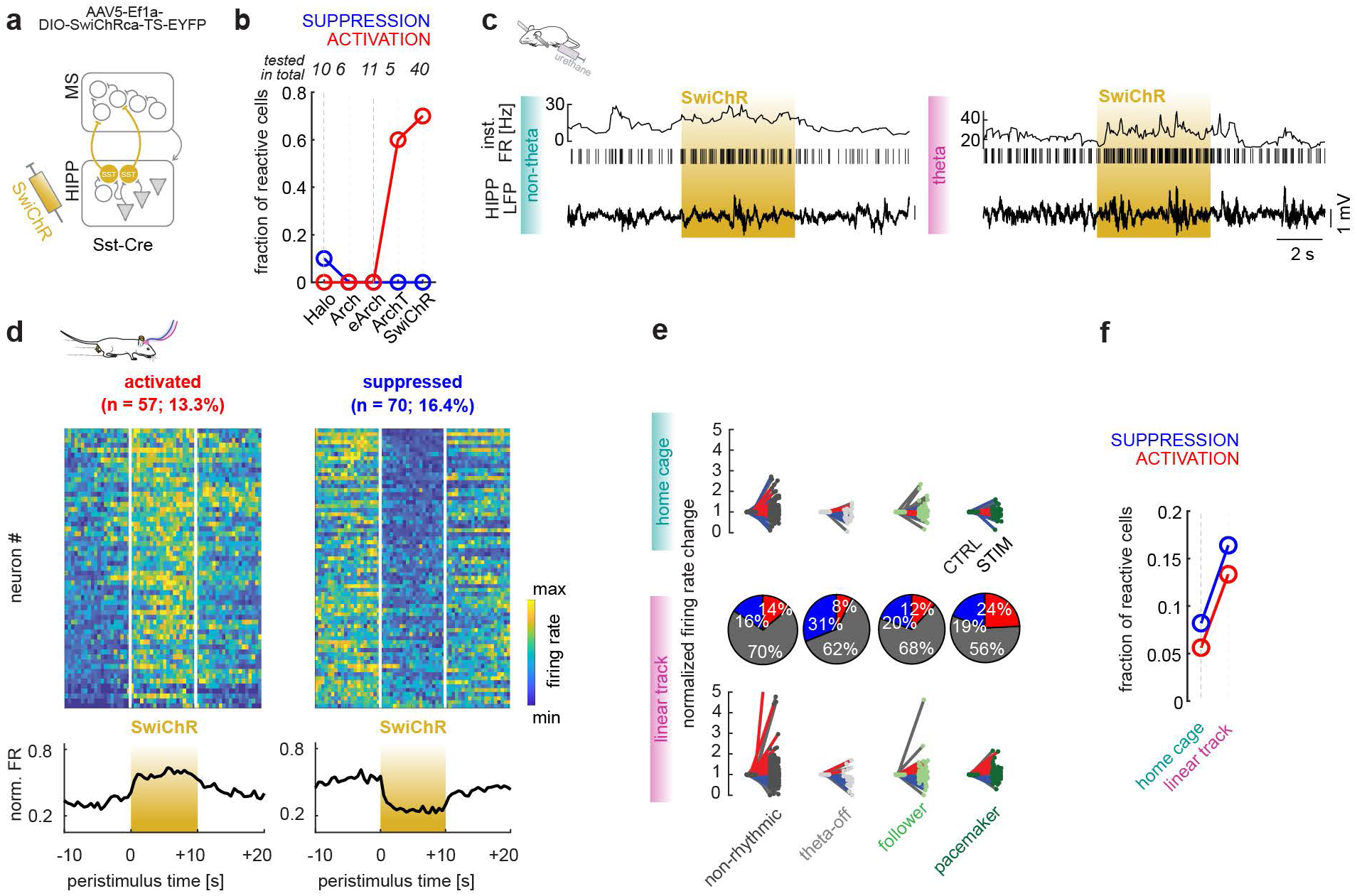
Optogenetic inhibition of HS feedback. **a,** Experimental design for optogenetic inhibition of the HS pathway in Sst-Cre mice injected in the hippocampus with AAV5-Ef1a-DIO-SwiChRca-TS-EYFP. Juxtacellular recordings under urethane anesthesia and chronic recordings in freely behaving animals were performed as in ChR2 experiments; in some juxtacellular recordings, Halo, Arch, eArch or ArchT were used instead of SwiChR (see respective Results section). **b,** Fraction of activated and suppressed MS neurons across inhibitory opsin experiments, under anesthesia. **c,** Effect of SwiChR mediated HS blockade on MS neuron firing under urethane anesthesia; illumination periods are marked by shaded boxes. **d**, Effect of HS blockade on MS firing in free behavior, in linear track. The normalized firing rates of significantly activated (left) or suppressed (right) neurons are shown. Bottom, population average. **e,** Top, firing rate change relative to control, in home cage, across rhythmicity groups during HS blockade. Middle, proportion of suppressed (blue) and activated (red) neurons in linear track. Bottom, firing rate change relative to control, in linear track. **f,** Fraction of activated and suppressed MS neurons in free behavior.

### Optogenetic activation of the HS pathway

MS neurons were grouped into four classes based on their firing autocorrelograms in home cage (or non-theta state) and linear track (or theta state) recordings using k-means clustering: the persistently theta-rhythmic pacemakers, the followers firing in theta-rhythm under hippocampal theta state, the theta-off cells that were non-rhythmic during theta but theta-rhythmic during non-theta and the non- rhythmics lacking theta-rhythmicity (Extended Data Fig. 3, Extended Data Tables 1,2). Among juxtacellularly labeled neurons, all tested pacemakers (6/6) and 1 out of 6 followers were identified as PV+ (Fig. 3d, Extended Data Table 1), corroborating earlier findings^8,45^. Neurons in all rhythmicity classes were inhibited upon ChR2-mediated activation of the HS pathway (Fig. 3f,g). Pacemaker neurons exhibited shorter-latency inhibition and more pronounced rebound firing (Fig. 3f,g, Extended Data Fig. 4), which in some cases resulted in a net increase in firing during the 10 s long stimulation trains (Extended Data Fig. 4l,n). Therefore, both suppression and activation during the whole stimulus train were observed among MS neurons, in both anesthesia and free behavior (Fig. 3h,i, Extended Data Fig. 4m,o), although activation was less prevalent in freely behaving animals (Fig. 3j,k).

Increasing stimulation frequency progressively enhanced both the firing rate change (Fig. 3k) and the fraction of responsive neurons (Extended Data Fig. 4p). The firing suppression was more pronounced in home cage (quiet periods) compared to linear track (exploration, locomotion; Fig. 3j,k), consistent with elevated baseline HS activity during movement inferred from our calcium imaging data (Fig. 1a-d). These findings indicate that HS pathway broadly inhibits MS neurons, with particularly strong effects on septal pacemaker cells. The graded modulation observed with increasing stimulation further supports that HS neurons transmit hippocampal population activity to MS circuits in a scalable manner.

### Optogenetic inhibition of the HS pathway

Optogenetic inhibition (Fig. 4a) provides a fast, reversible loss-of-function test of the function of the HS pathway, expected to disinhibit postsynaptic MS neurons (Fig. 4b), and reveal the contribution of the HS feedback to theta genesis. In anesthetized animals, multiple inhibitory opsins (Halo, Arch, eArch, ArchT and SwiChR) were evaluated using juxtacellular MS recordings and local optogenetic illumination (Fig. 4b). SwiChR elicited the most robust effects (Fig. 4b,c) and was therefore used in freely behaving experiments (Fig. 4d,e). Consistent with higher baseline HS activity during locomotion (Fig. 1a-d), SwiChR-mediated effects were more prominent on the linear track than in the home cage (Fig. 4f).

SwiChR-evoked blockade of the inhibitory HS pathway led both to activation and suppression of a comparable fraction of MS neurons. In freely behaving mice, pacemakers were activated in the highest proportion (Fig. 4d,e). Activation of followers and pacemakers during HS suppression was accompanied by increased bursting propensity (as reflected by higher burst index and burst occurrence^46^ (Extended Data Fig. 5b,d), whereas suppressed pacemakers exhibited reduced bursting (Extended Data Fig. 5b,d). Intra-burst frequency and burst length, reflecting intrinsic burst dynamics^47^ were only minimally affected (Extended Data Fig. 5f,h). Complementary results were observed in ChR2 experiments, where the suppressed rhythmic neurons exhibited reduced bursting (Extended Data Fig. 5a,c) with minimal changes in intra-burst frequency and burst length (Extended Data Fig. 5e,g). These results indicate that HS input modulates the bursting propensity of MS neurons but is not required for burst generation, while intrinsic burst dynamics remain largely preserved.

### Rhythmicity and synchrony of MS neurons in response to manipulation of the HS feedback

Beyond changes in firing rate and burst properties, HS feedback may also alter the rhythmic firing of individual MS neurons or their synchrony. Theta rhythmic firing was quantified by theta modulation depth, which was reduced in suppressed MS neurons in HS-activation experiments (Fig. 5a). Complementing this, HS blockade by SwiChR augmented theta rhythmicity of activated pacemakers (Fig. 5b). Although in the case of some pacemakers with no firing rate change, increased rhythmicity upon illumination was observed, on the group level, unaffected or suppressed pacemakers showed no rhythmicity change upon HS blockade (Fig. 5b).

**Figure 5.**
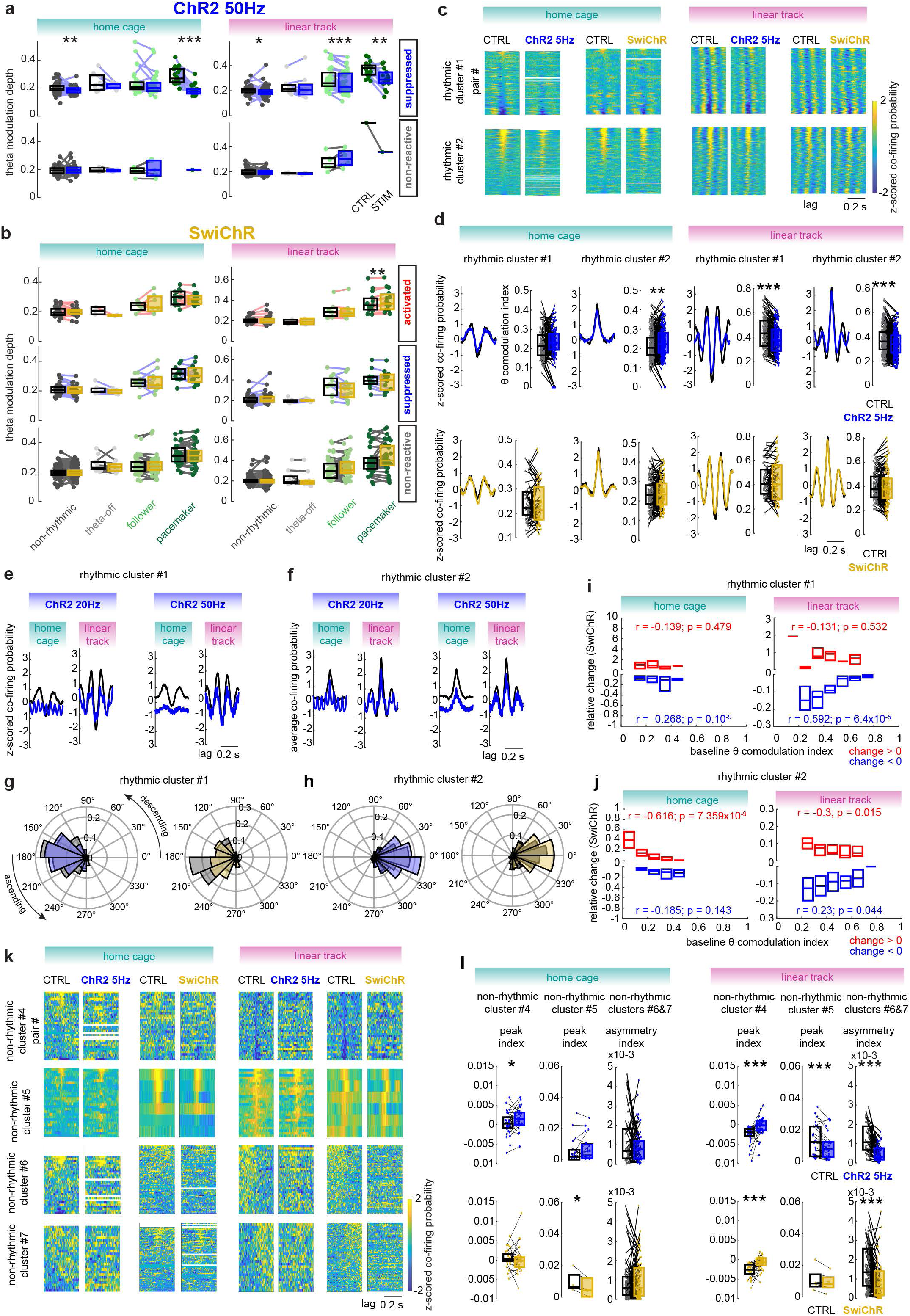
Synchrony of MS neurons. **a-b,** Theta modulation depth of individual MS neurons during ChR2 50 Hz (a) and SwiChR (b) stimulation in freely behaving mice. Wilcoxon signed-rank tests (control vs stimulation): a, home cage (HC), suppressed, non-rhythmics (N), p = 0.0025, theta-offs (O), p = 0.0625, followers (F), p = 0.3217, pacemakers (P), p = 0.000196; non-reactive, N, p = 0.8613, O, p = 0.75, F, p = 0.4375; linear track (LT), suppressed, N, p = 0.0263, O, p = 0.9453, F, p = 0.000163, P, p = 0.00434; non-reactive, N, p = 0.674, O, p = 1, F, p = 0.0625, P, p = 1; b, HC, activated, N, p = 0.5435, O, p = 0.5, F, p = 0.5781, P, p = 0.2769; suppressed, N, p = 0.1526, O, p = 0.4688, F, p = 0.6377, P, p = 0.5186; non-reactive, N, p = 0.294, O, p = 0.5772, F, p = 0.4688, P, p = 0.3172; LT, activated, N, p = 0.86, O, p = 1, F, p = 0.9375, P, p = 0.00427; suppressed, N, p = 0.1303, O, p = 0.2188, F, p = 0.2334, P, p = 0.1514; non-reactive, N, p = 0.2593, O, p = 0.625, F, p = 0.3542, P, p = 0.1218; **c,** Color-coded cross-correlograms (CCGs) of rhythmic clusters #1 and #2. **d,** Population average CCGs and the theta comodulation index (control versus stimulation) of rhythmic clusters #1 and #2. Boxes show median and interquartile range. Wilcoxon signed-rank tests: cluster #1, ChR2 5Hz, HC, p = 0.12756, LT, p = 1.4887×10^-11^; SwiChR, HC, p = 0.25414, LT, p = 0.17261; cluster#2, ChR2 5 Hz, HC, p = 0.0017333, LT, p = 2.1749×10^-7^; SwiChR, HC, p = 0.52272, LT, p = 0.13156. **e-f,** Population-average CCGs of pairs in rhythmic cluster #1 (e) and #2 (f) during 20 Hz (left) and 50 Hz (right) ChR2 stimulation in home cage and linear track. **g-h,** CCG phase at zero lag of pairs in rhythmic cluster #1 (g) and #2 (h) during 5 Hz ChR2 (left) and HS blockade (right). **i-j,** Change in theta comodulation index relative to control in rhythmic cluster #1 (i) and #2 (j) during HS blockade, binned by baseline theta comodulation index; increases and decreases shown separately. Spearman’s ρ and p-values are shown for the correlation between control comodulation and change, calculated separately for pairs with increase or decrease, data were unbinned. **k,** Color-coded cross-correlograms (CCGs) of non-rhythmic clusters #4-7. **l,** Peak index for non-rhythmic cluster #4-5 and asymmetry index for non-rhythmic clusters #6&7 (pooled, absolute values used) during 5 Hz ChR2 and SwiChR stimulation. Wilcoxon signed-rank tests: cluster #4, ChR2 5 HZ, HC, p = 0.035009, LT, p = 1.9938×10^-5^; SwiChR, HC, p = 0.097772, LT, p = 2.5967×10^-5^; cluster #5, ChR2 5 Hz, HC, p = 0.34088 (n = 10 outliers not shown in the plot), LT, p = 0.00024254; SwiChR, HC, p = 0.03125 (n = 1 outlier not shown), LT, p = 0.09375; clusters #6&7, ChR2 5 Hz, HC, p = 0.83558 (n = 1 outlier not shown), LT, p = 6.963×10^-11^ (n = 3 outliers not shown); SwiChR, HC, p = 0.41167 (n = 10 outliers not shown), LT, p = 4.0261×10^-9^ (n = 17 outliers not shown).

Crosscorrelograms (CCGs) were computed to examine the synchrony between the simultaneously recorded MS neuron pairs in freely behaving mice. Based on the CCGs, we identified rhythmic and non-rhythmic clusters of neuronal pairs (Extended Data Fig. 6a-c). Rhythmic clusters were composed mainly of pairs of rhythmic (follower or pacemaker) neurons (Extended Data Fig. 6d), and these pairs exhibited strong comodulated firing (anti-phase comodulation in cluster #1 and in-phase comodulation in cluster #2) at theta rhythm (quantified as theta comodulation index) in linear track (Fig. 5c, Extended Data Fig. 6c), and weak comodulation in home cage (Extended Data Fig. 6a,c). HS activation by ChR2, which strongly reduced overall firing rates (Fig. 3i,j) and theta modulated firing (Fig. 5a), also dampened theta comodulation of rhythmic neuron pairs (Fig. 5c-f). However, the coupling phase was preserved on the linear track (Fig. 5d,g,h), even at 20 or 50 Hz ChR2 stimulation (Fig. 5.e,f). In home cage, by contrast, the weaker baseline synchrony between MS neurons was less resistant to the modulation by stimulation (Fig. 5e,f). During SwiChR-mediated HS blockade, comodulation depth and phase of rhythmic pairs were largely preserved (Fig. 5c,d), but - surprisingly - pairs with the smallest comodulation index were more affected, whereas pairs with stronger rhythmic coupling were more resilient to HS suppression (Fig. 5i,j). These findings underscore the dominance of intra-MS circuits over the HS feedback in theta rhythm generation, consistent with our assumptions from the *in silico* modeling. HS feedback rather suppresses the MS theta generator than it is required for generating the rhythm.

In addition to the theta rhythmic neuron pair clusters, we identified non-rhythmic clusters with negative or positive coupling (clusters #4 and #5), as well as asymmetric, ramping-like coactivity patterns (clusters #6 and #7; Extended Data Fig. 6a,c). These coupling types – excluding cluster #5 with prominent zero lag co-firing - were robustly reduced by optogenetic manipulations of the HS pathway in the linear track, using both ChR2 and SwiChR (Fig. 5k,l). Notably, SwiChR manipulation in linear track shifted non-rhythmic coactivity toward a pattern resembling that observed during control home cage recordings. In contrast, SwiChR (and ChR2) manipulation had only marginal effects on non-rhythmic coactivity in the home cage. These results further indicate that intra-MS connections among non-rhythmic neurons (likely generating non-rhythmic CCGs, Extended Data Fig. 6d) are more sensitive to the HS feedback in a behavioral-state-dependent manner than the well-established rhythmic interactions among rhythmic neurons.

### Hippocampal effects of HS manipulations

We next tested whether coupling to hippocampal oscillations changed during optogenetic manipulation of the HS feedback, beginning with an analysis of hippocampal LFPs. Stimulation trains drove the hippocampal pyramidal-layer local field potential at the stimulation frequency (Fig. 6a, Extended Data Fig. 7g,i), whereas theta score (relative theta power) on the linear track was largely unchanged (Fig. 6c,e Extended Data Fig. 7e). Five Hz stimulation, overlapping with the intrinsic theta range, caused a small increase of theta score on the linear track (Fig. 6c,e). HS blockade produced a modest but highly significant increase in the ongoing hippocampal theta power on the linear track (Fig. 6b,d,e, Extended Data Fig. 7a,c,d,e). Because stimulations were delivered pseudo-randomly, this theta increase could, in principle, reflect higher running speed during stimulation epochs. However, animals did not run significantly faster during SwiChR trials (Fig. 6b, Fig. 7l), and comparing theta scores within matched speed bins did not abolish the significant theta increase (Fig. 6f,g). Moreover, the length of theta cycles was unchanged by HS blockade (Fig. 6h). Consistent with the increase in theta score during 5 Hz ChR2 stimulation, speed-corrected spectral changes showed elevated power across speed bins (Extended Data Fig. 7f), whereas during HS blockade, the speed-corrected theta increase was restricted to bins corresponding to active movement (Fig. 6g).

**Figure 6.**
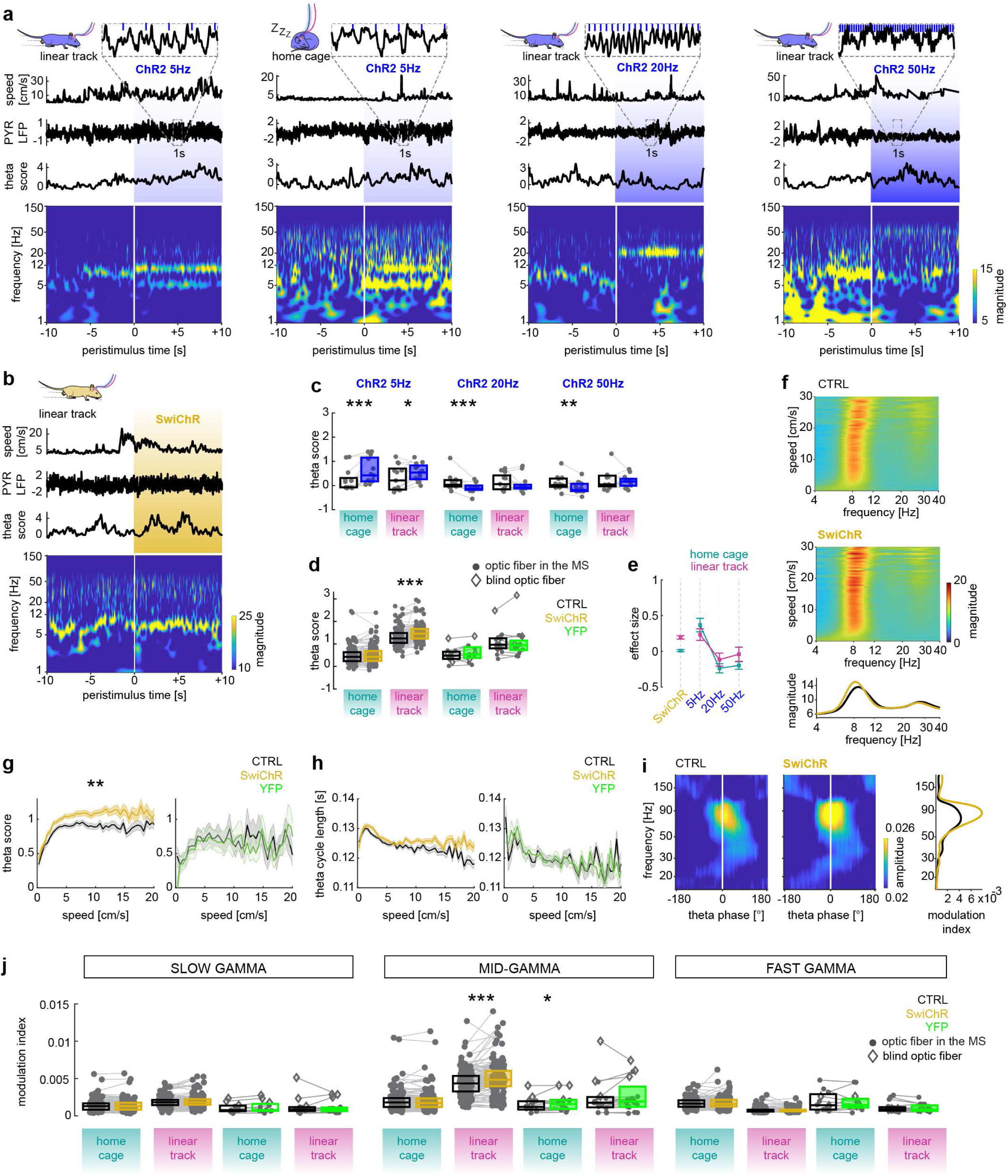
Hippocampal effects of HS manipulations. **a-b,** Examples of ChR2 stimulations at 5 Hz in linear track, 5 Hz in home cage, 20 Hz in linear track, 50 Hz in linear track (a, left to right) and SwiChR stimulation in linear track (b). From top to bottom: animal speed (3D head position), z-scored CA1 pyramidal layer LFP (insets show zoomed periods of examples, blue ticks indicate light pulses during stimulation), theta score ((theta magnitude – delta magnitude)/delta magnitude), and LFP spectrogram (white line: stimulation onset, shaded area: stimulation). **c,** Theta score during ChR2 stimulation at 5, 20 and 50Hz (left to right, n = 13 for each). Wilcoxon signed-rank tests: 5Hz home cage (HC), p = 0.00048828; 5Hz linear track (LT), p = 0.01709; 20Hz HC, p = 0.00024414; 20Hz LT, p = 0.6355; 50Hz HC, p = 0.003418; 50Hz LT, p = 0.94604. **d,** Theta scores during HS blockade (n = 126 recording sessions) versus YFP controls (home cage, n = 12, linear track, n = 16 sessions) in the home cage and on the linear track. HC, two-way RM ANOVA (stimulation x opsin): F(1,136) = 2.3151, p = 0.1304. LT, two-way RM ANOVA (stimulation x opsin): F(1,140) = 7.7882, p = 0.006; multiple comparison (control vs stimulation) within groups: HC SwiChR, p = 0.5207, HC YFP, p = 0.0734; LT SwiChR, p = 1.0597×10^-10^, LT YFP, p = 0.9601. Dot markers indicate illumination with implanted optic fibers; diamonds indicate fibers above the skull (blind fibers). **e,** Effect size of stimulation on the theta score (stim-ctrl); error bars indicate mean ±SEM. **f,** Frequency spectra (magnitude from wavelet decomposition) across binned speeds, control (top) and SwiChR mediated HS blockade sessions (middle). Bottom, speed-averaged spectra highlighting enhanced power around 8 Hz. **g-h,** Theta score (l) and theta cycle length (m) across speed bins in SwiChR and YFP recording sessions; shaded areas indicate mean ±SEM. For theta score, two-way RM ANOVA (stimulation x opsin): F(1,134) = 7.4541, p = 0.0072; three-way RM ANOVA (stimulation x opsin x speed): F(1,134) = 6.6985, p = 0.0107. For theta cycle length, two-way RM ANOVA (stimulation x opsin): F(1,134) = 2.2803, p = 0.1334; three-way RM ANOVA (stimulation x opsin x speed): F(1,134) = 3.0306, p = 0.084. Note that some sessions lack data at higher speed bins because the animal did not reach those speeds; ANOVAs were restricted to 5-10 cm/s bins. **i,** Cross-frequency coupling between CA1 pyramidal layer theta phase and supra-theta amplitude in an example linear track recording session. Phase-amplitude plots for control (left) and HS blockade (middle); white line marks theta peak (0° phase). Right, modulation index in control (black) and HS blockade (yellow), showing increased coupling in the 60-90 Hz band. **j,** Modulation index of CA1 pyramidal layer cross-frequency coupling for slow (40-60Hz), mid (60-90Hz), and fast (120-150Hz) gamma ranges in SwiChR experiments. Asterisks indicate the significant differences revealed by multiple comparison (control vs stimulation). Two-way RM ANOVA (stimulation x opsin) tests: slow gamma in HC, F(1,136) = 1.5748, p = 0.2117; multiple comparison within SwiChR, p = 0.2657, within YFP, p = 0.0976; slow gamma in LT, F(1,140) = 1.0222, p = 0.3137; multiple comparison within SwiChR, p = 0.0526, within YFP, p = 0.702; mid-gamma in HC, F(1,136) = 4.6416, p = 0.033, multiple comparison within SwiChR, p = 0.8692, within YFP, p = 0.0275; mid-gamma in LT, F(1,140) = 1.627, p = 0.2042, multiple comparison within SwiChR, p = 1.1148×10^-9^, within YFP, p = 0.4109; fast gamma in HC, F(1,136) = 0.0505, p = 0.8226, multiple comparison within SwiChR, p = 0.4889, within YFP, p = 0.9828; fast gamma in LT, F(1,140) = 1.4308, p = 0.2337, multiple comparison within SwiChR, p = 0.6124, within YFP, p = 0.147.

**Figure 7.**
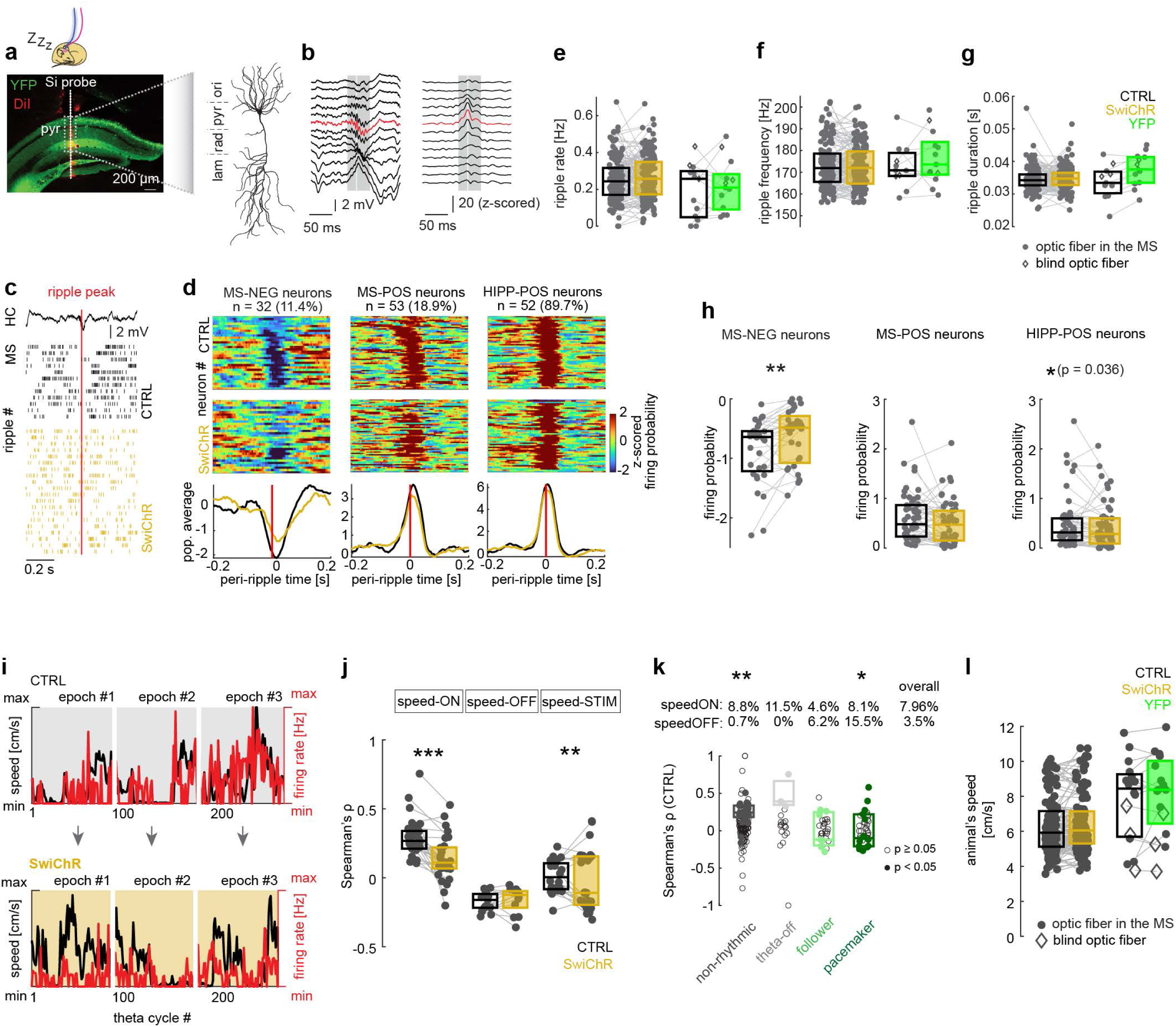
Effect of manipulating HS feedback on MS neurons’ ripple- and movement speed coupling. **a,** Histological reconstruction of the probe track (white line, dots indicating recording sites; the probe was DiI-coated prior surgery) implanted into hippocampus for chronic recordings in free behavior, with a schematic CA1 pyramidal neuron (scidraw.io) for recording site locations along the probe. **b,** Example raw (left) and ripple band (150-250 Hz) magnitude of hippocampal LFP recorded in freely behaving animal in home cage; the channel with maximal ripple power is shown in red. White line indicates ripple peak, grey shading the ripple duration. **c,** Firing of an example MS neuron aligned to hippocampal ripple peaks in control and SwiChR mediated blockade of HS terminals in the MS. **d,** Firing probability of negatively modulated (MS-NEG) or positively modulated (MS-POS or HIPP-POS) neurons related to hippocampal ripple peaks in control and SwiChR mediated HS blockade. Note that hippocampal (HIPP) neurons were also tested. Top, individual neuronal data; bottom, population average. **e-g**, Ripple rate (e), ripple frequency (f) and duration (g) upon HS blockade, compared to controls and recordings from YFP animals. **h,** Firing probability around ripple peaks in control and HS blockade; Wilcoxon signed-rank tests (control versus stimulation): MS-NEG, p = 0.0012989; MS-POS, p = 0.05417; HIPP-POS, p = 0.036207. **i,** Firing rate (red) of an example MS neuron and the simultaneously registered speed of the animal (black), both computed as per-theta-cycle averages. White lines delineate demonstrative 10 s control (preceding, upper row) and stimulation epochs (bottom); inter-epoch intervals varied (not shown), and theta cycle counts differed between epochs. Note the reduced speed-firing correlation during HS blockade. **j,** Spearman’s ρ between per-theta-cycle speed and firing rate (included cycles 50-250 ms and firing rate > 0Hz) for control-identified speed-ON (ρ > 0, p < 0.05; n = 34) speed-OFF (ρ < 0, p < 0.05; n = 15) and speed-STIM MS neurons (p< 0.05 only during stimulation, irrespective of sign of ρ; n = 28; see Methods for criteria). Wilcoxon signed-rank tests (SwiChR vs control): speed-ON, p = 8.8479×10^-6^; speed-OFF, p = 0.41431; speed-STIM, p = 0.0015073 (in this case |ρ|). **k,** Fraction of speed-correlated MS neurons across rhythmicity groups. Box plots show control Spearman’s ρ distributions only for significant speed-ON/OFF neurons (empty dots: non-significant neurons); population fractions are indicated above columns per rhythmicity class and overall. Rhythmicity vs speed category: χ²(3) = 17.13, p = 0.001, Fisher’s post-hoc test (Bonferroni-corrected): non-rhythmics, p = 0.002, theta-offs, p = 1, followers, p = 0.718, pacemakers, p = 0.017. **l,** Per-theta-cycle speeds across all recording sessions (n = 126 SwiChR, n = 16 YFP control sessions). Two-way RM ANOVA (stimulation x opsin): F(1,140) = 0.7010, p = 0.4039; multiple comparison tests for control vs stimulation within the SwiChR or YFP sessions: p = 0.2226 (SwiChR), p = 0.1857 (YFP).

In addition, theta increase during HS blockade was accompanied by a significant enhancement of mid-gamma oscillations (of likely entorhinal origin^48^) nested within theta cycles (Fig. 6i,j). These results indicate that augmented theta upon HS feedback suppression may reflect elevated synchrony throughout the limbic network.

MS neurons across all rhythmicity groups and ChR2 stimulus conditions became locked to the stimulation-entrained hippocampal oscillations (Extended Data Fig. 7g-j), indicating that the stimulation rhythm acted as a common driver for MS and hippocampus, possibly by retrogradely activating HS neurons and thereby pacing hippocampal circuits via their local efferents. Under urethane anesthesia, we observed similar hippocampal LFP changes during ChR2 stimulation (Extended Data Fig. 7a,b,d), but no significant changes during HS blockade (Extended Data Fig. 6c,d). MS neurons were also strongly locked to the stimulation-entrained hippocampal rhythm under urethane (Extended Data Fig. 7g,h). Because ChR2 stimulation entrained hippocampal oscillations leading to artificial phase-locking, theta phase preference of MS neurons was evaluated in SwiChR experiments only.

During HS blockade, the MS followers and pacemakers preserved their preferred theta phase (Extended Data Fig. 7k), but the phase locking of pacemakers became slightly yet significantly stronger (Extended Data Fig. 7l). Hippocampal neurons preserved their coupling strength to theta during HS blockade (Extended Data Fig. 7m). Together with CCG observations where the coupling phase was unaltered during HS suppression (Fig. 5g,h), these data suggest that HS feedback is not required for maintaining synchrony among MS pacemakers nor for their phase alignment to hippocampal theta.

### Ripple-locked firing of MS neurons during blockade of the HS feedback

In addition to the observation of ripple-coupled HS neurons in vitro (Fig. 1k-n), we also identified negatively and positively ripple-locked MS neurons in freely behaving mice (Fig. 7c,d, resembling to earlier data^49^). Although SwiChR mediated HS blockade preserved ripple rate, frequency and duration (Fig. 7a,b,e-g), the ripple locking of negatively coupled MS neurons was reduced (Fig. 7c,d,h). Ripple-locked activation of MS neurons and the coupling of hippocampal neurons were only minimally affected (Fig. 7d,h). According to this finding, a large part of the MS circuit is transiently suppressed during ripples by pyramidal cell-recruited HS feedback (Fig. 1k-n).

### HS blockade reduces the speed-modulated firing of MS neurons

Among MS neurons, we identified a subpopulation with speed-modulated firing (Fig. 7i,j). Positively speed-modulated neurons (speed-ONs) were found predominantly in the non-rhythmic group, but, interestingly, also among pacemakers which contained most of the speed-OFF cells, whose firing was anticorrelated with speed (Fig. 7k). During HS blockade, speed modulation of speed-ON cells was reduced (Fig. 7j), while in some neurons not exhibiting speed-modulation in control periods, HS suppression led to the emergence of speed-modulated firing (speed-STIM cells, Fig. 7j). These observations indicate that the movement-tuned HS feedback (see also the Ca-imaging results) may be an overlooked source of speed-related activity of MS neurons conveying a speed signal, possibly originating from the hippocampal principal cell populations^50^ (Fig. 1a-d). Thus, the HS feedback exerts multifaceted regulation of MS neurons, but is not required for theta genesis as previously was broadly assumed.

## DISCUSSION

Long-range GABAergic neurons are a distinct and still relatively poorly understood component of cortical and hippocampal circuits^27,44,51^. Their function is suggested to synchronize oscillations, as outlined in case of hippocampal-entorhinal reciprocal inhibitory projections^52,53^, or to exert phase control, as the HS projection may regulate and facilitate theta-rhythmic spiking in the MS^27,29,49^. A related model of theta generation^24^ posits that HS neurons are required elements of an alternating inhibitory–disinhibitory loop for theta rhythmogenesis. Understanding the generation and coordination of theta across limbic structures is therefore essential for deciphering the fundamental mechanisms of how various information streams are orchestrated during episodic memory encoding and retrieval^54,55^. Here we address this question by characterizing the behavior-dependent recruitment of the HS pathway and its impact on MS activity.

Guided by the studies above, we hypothesized that selective inhibition of the HS projection would reduce theta oscillations throughout the limbic system by attenuating the theta bursting of MS neurons, leading to the suppression of hippocampal theta oscillations and to the disruption of septo-hippocampal phase-coupling. To achieve selective HS manipulations, we devised the opsin-mediated suppression of the MS-targeting projection of SST-positive hippocampal neurons.

Contrary to our expectation, optogenetic suppression of HS pathway preserved septal and hippocampal theta, and in freely behaving animals, theta even became more prominent. The strong impact of HS blockade on MS neuron activity and LFP on a linear track is consistent with Ca-imaging in head-restrained preparations, where HS neurons show elevated activity during movement. Although optogenetic inhibition of axon terminals is inherently incomplete, our data suggest that HS feedback is not required for theta genesis, despite being capable of rhythmically entraining septo-hippocampal circuits as seen in the ChR2-mediated HS activation. However, in this latter case, antidromic activation of HS neurons^56^ and the MS-independent recruitment of the hippocampus because of the extensive hippocampal axon arbors of HS neurons^33^ cannot be excluded. Thus, SwiChR-mediated HS blockade was employed to resolve the central tension of our study: HS neurons act as powerful modulators of MS activity yet are dispensable for theta genesis.

Suppression of HS input increased theta-modulated bursting in pacemaker MS neurons in a state-dependent manner, consistent with higher baseline HS drive during ongoing hippocampal theta. Theta-modulated coupling between MS neuron pairs was unexpectedly resilient to both SwiChR-mediated HS blockade and ChR2-mediated HS activation, even when MS firing rates were strongly reduced during ChR2 50 Hz stimulation. Theta modulation strength of single neurons and phase relationships among pacemakers and followers also remained remarkably stable. Moreover, pairs with stronger theta comodulation were more resistant to HS perturbations, suggesting that established intra-MS coactivity becomes relatively independent of hippocampal control, whereas more weakly coupled pairs remain susceptible to top-down modulation - consistent with the view that theta is primarily generated within the MS and then propagates to the hippocampus^9,19^. Overall, our results support our current MS model in which intra-MS circuitry dominates over HS feedback in theta genesis. The bidirectional changes in MS firing observed during HS blockade and activation may arise from an inhibition-stabilized MS network, where perturbation of a single (HS) input induces compensatory adjustments across local GABAergic (and cholinergic, glutamatergic) populations.

Modulation of responsive MS rhythm-generating neurons can tune theta power and adapt it to hippocampal population activity, revealing a novel feedback-control circuit motif in the episodic memory system. In line with this concept, selective blockade of the HS projection during movement increased hippocampal theta power, an effect that remained robust after correcting for running speed^57^. Thus, the HS projection inhibits theta rhythmogenesis. The functional relevance of this effect is further supported by parallel changes in theta-coupled higher-frequency activity (mid-gamma), which reflects entorhinal–hippocampal communication^48^. These coordinated shifts indicate that optogenetic manipulation reshaped a distributed network involving multiple brain regions rather than inducing a purely local perturbation. Consistent with the hippocampal integration of subcortical modulatory signaling and external salient events, HS neurons were indirectly affected by cholinergic modulation.

Besides theta, ripples are a prominent hippocampal discharge pattern with a key role in memory processing^58^. HS neurons - likely a specific subpopulation when considering our data together with earlier findings^35^ - are temporally aligned to hippocampal ripples, implicating them in ripple control. Although basic ripple characteristics were unchanged by HS feedback inhibition, ripple-related suppression of MS neurons was attenuated, pointing to a modulatory role of HS neurons during ripples, analogous to their modulatory influence during theta. The HS feedback is well positioned to dynamically tune septal circuits, including non-rhythmic neurons^49^. To fulfill this role, HS neurons faithfully track hippocampal network activity and relay it in a graded, state-dependent manner: activating their numerous excitatory inputs over a wide range elicits quantitatively coherent, quasi-linear changes in firing, and their axons can modulate MS activity in a firing-rate-dependent manner. When further speculating about the function of the pyramidal cell-controlled HS feedback, we should consider the common mechanism behind elevated pyramidal cell activity and positive movement-correlation of theta power and frequency during exploration: higher impulse flow from brainstem and hypothalamic activating systems including subcortical modulatory pathways. According to our results, the HS feedback may be key for compensating this ascending, theta-driving excitation at the level of the main theta generator, the MS. Moreover, the movement-associated inhibition by the HS feedback may also have the unexpected effect of rendering MS neuronal activity speed-correlated.

Corroborating earlier findings from slice experiments^29^, ChR2-mediated activation of HS fibers elicited a robust rebound increase in the firing of pacemaker-type MS neurons, most prominently during theta in the linear track. This rebound emphasizes the contribution of intra-MS circuits, where local synaptic connectivity and intrinsic membrane properties^8^ compensate transient inhibitory HS input and help preserve pacemaker timing. During theta, these mechanisms are particularly effective (whereas dampened in urethane, resulting in weaker rebound and less HS-blockade–induced inhibition of MS firing), making the rebound more pronounced. The reduction of MS firing during HS suppression further reveals intra-MS mechanisms: disinhibited neurons exert stronger inhibition onto others that receive weaker HS input. Consistent with this assumption, our MS model supports a dominant role for intra-MS connectivity in establishing and sustaining theta, even though strong HS drive can transiently override ongoing rhythms and impose the stimulation frequency on both MS and hippocampal activity, in agreement with our and others^30^ experimental observations. This bidirectional, population-level modulation offers a simple explanation for why hippocampal theta power shows only modest overall changes - tending to increase rather than decrease - despite robust alterations in MS firing, and it helps reconcile long-standing mixed findings^59^ on MS contributions to hippocampus-dependent behavior when non-selective manipulations recruit multiple MS cell types simultaneously.

Because the dorsal CA1 HS population is heterogeneous and dispersed, and *in vivo* recordings from identified HS neurons during behavior remain technically challenging, we focused on the postsynaptic side and quantified how MS neurons respond to HS manipulation. Back-labeling from the MS likely transduced not only SST-positive HS interneurons but also some pyramidal cells projecting to lateral septum or directly to MS^38^. Future work employing more precise anatomical and genetic targeting of HS subpopulations—including MS-projecting principal cells—will be required to dissect how distinct HS subnetworks are recruited and how they differentially shape septal and hippocampal dynamics. Our conclusions rely largely on optogenetic inhibition of HS feedback. However, because the manipulation was likely incomplete, residual HS feedback may have been sufficient to sustain theta generation, potentially explaining the relatively subtle changes observed in theta-rhythm-related firing of MS neurons. Contradicting this possibility, the same manipulation robustly affected coactivity among non-rhythmic MS neurons, as well as ripple-related and speed-modulated firing of MS neurons. We therefore consider it highly unlikely that the incompleteness of HS blockade accounts for our observations.

Notably, additional MS inputs, e.g. from the supramammillary nucleus and brainstem, may counteract HS top-down effects, reducing the observed changes in our experiments. Finally, we may also consider additional functions unfolding over a longer timescale, beyond the fast modulation of MS activity, which are not studied here. As demonstrated previously^60–62^, longer-lasting effects of the feedback conveyed by somatostatin targets slow-firing, putative cholinergic neurons and release of nerve growth factor would translate hippocampal activity states into durable plastic changes of the MS circuit.

## ACKNOWLEDGEMENTS

We express our gratitude for the excellent technical assistance by Katalin Lengyel and for the administrative help by Katalin Ivanyi. We also thank Thomas R. Reardon for providing the rabies and pseudorabies constructs. We also thank Andre Berndt and Soo Yeun Lee for SwiChR. Mouse drawings and schematic pyramidal cell were downloaded from SciDraw.

## AUTHOR CONTRIBUTIONS

A.D., P.H. M.J., L.L., E.P., D.S., G.T., V.V. performed the experiments; B.K. carried out network simulations; A.D., D.S., G.T. analyzed the electrophysiology and imaging data; G.N. and R.K. analyzed anatomical experiments; K.D. provided optogenetic constructs; A.I.G., B.H., A.L., V.V. supervised experiments. T.F.F., G.N., V.V. acquired funding. A.D., B.K., D.S. and V.V. wrote the paper with significant input from B.H. and M.J. and feedback from all authors; V.V. supervised the project.

## FUNDING

This work was supported by NKFIH K109790 and K132735 to VV., ERC-AdG SERRACO to T.F.F., NKFIH K115441 and KH124345 to AIG., a Postdoctoral Fellowship (R483-2024-1854) from the Lundbeck Foundation to D.S., Frontline Research Excellence Program of the Hungarian National Research, Development and Innovation Office, NRDI Fund 133837 (to GN), Hungarian Brain Research Program, NAP3.0 NAP2022-I-1/2022 (to GN). The funders had no role in study design, data collection, analysis and publishing.

## METHODS

### HS cell functional imaging Animals

HS cell functional imaging experiments were conducted at the Department of Neurosciences, Columbia University, in accordance with the US National Institutes of Health guidelines and with approval from the Institutional Animal Care and Use Committees of Columbia University and the New York State Psychiatric Institute. We used wild-type C57BL/6 mice and hemizygous offspring of the Pvalb-Cre mice crossed with the Ai9 reporter line on a C57BL/6 background (Ai9: loxP-STOP-loxP-tdTomato Cre reporter strain B6;129S6-Gt(ROSA)26Sor^tm9(CAG-tdTomato)Hze/J^, Jackson Laboratory)

### Injections and surgery

The MS was injected with either CVS-N2c^ΔG^-GCaMP6f (wild-type mice, n = 5) or the EnvA-pseudotyped CVS-N2c^ΔG^-GCaMP6f together with rAAV1/2(CAG-TVA-mCherry)Cre and rAAV1/2[EF1a-histoneGFP-2A-rabiesG(N2c)]Cre helper viruses (Pvalb-Cre mice, n = 3) to retrogradely label the HS cells projecting to the MS^36^. For hippocampal window implantation, mice were anesthetized with isoflurane and received buprenorphine (0.1 mg/kg, s.c.) for perioperative analgesia. Mice were head-fixed in a stereotaxic frame, the skull was exposed, and a 3-mm diameter craniectomy was made above the left dorsal CA1 region, matching the diameter of the cannula window.

After removing the dura, the overlying cortex above the hippocampus was slowly aspirated under continuous irrigation with chilled cortex buffer until the capsula externa was exposed. A sterilized cannula window was then gently wedged into place, and both the cannula and a stainless-steel head-post were secured to the skull with grip cement; adhesion to bone was strengthened using Kerr OptiBond dental adhesive. Mice were returned to their home cage after 15-20 min (cement curing time) and typically regained full mobility within 5-20 min. Animals were monitored every 12 h for 3 days post-surgery and received buprenorphine to minimize discomfort. Beginning 3 days after surgery, mice were habituated to handling and head-restraint, after which imaging of GCaMP6f-expressing hippocampal cells was initiated.

### Two-photon imaging

*In vivo* imaging was performed using a resonant galvo-based two-photon microscope (Bruker) equipped with an ultra-fast pulsed laser (Coherent; 920 nm) controlled by an electro-optical modulator and a 40× water-immersion objective. Distilled water was used to couple the objective to the cannula window. Fluorescence was collected through a GaAsP photomultiplier tube (green channel for GCaMP6f), operated with PrairieView software. Head-fixed mice ran on a linear treadmill during imaging. Goniometer was used to tilt the head up to 10° so that the imaging window was parallel to the objective.

Locomotor behavior was monitored by measuring treadmill wheel rotation, recorded as voltage changes from an infrared phototransistor as wheel spokes intermittently blocked light from an infrared LED. Analog signals encoding running speed were acquired with an analog-to-digital converter and synchronized to two-photon imaging by a common trigger pulse. Time series were collected in the green channel (GCaMP6f) at 512 × 512 pixels at ∼30 Hz and motion-corrected as described previously^63^.

### Data analysis

Regions of interest (ROIs) were manually drawn on motion-corrected time series in ImageJ (NIH) to isolate somata of HS cells, and ΔF/F calcium signals (∼30 Hz sampling rate) were extracted. Calcium traces were temporally aligned with running speed signals (sampled at 100 Hz) by interpolating to the imaging frame rate. Only data from n = 2 wild-type and n = 2 Pvalb-Cre mice were included in the final analysis. The somatostatin content of GCaMP6f-labelled hippocampal cells in one illustrative animal was confirmed by post hoc SST immunostaining: free-floating sections were washed three times in PBS (3 × 10 min) and then incubated with rat anti-somatostatin primary antibody (1:100, Millipore MAB354) diluted in antibody diluent overnight at 4°C on a shaker. Sections were then washed three times in PBS (3 × 10 min) and incubated with a species-specific secondary antibody (anti-rat secondary conjugated with Alexa594, 1:300 in antibody diluent) for 2 h to overnight at room temperature/4°C. After a final series of three PBS washes (3 × 10 min), sections were mounted on glass slides, air-dried, and coverslipped with mounting medium.

To quantify the relationship between locomotion and HS cell activity, Spearman’s rank correlation coefficient (ρ) was computed between running speed and calcium signal for each HS cell. Spearman’s ρ values were then Fisher z-transformed (z = 0.5ln((1+ ρ)/(1- ρ)), to stabilize the variance across correlation strengths. The mean Fisher-transformed value across cells was calculated and inverse-transformed to obtain a population-level correlation estimate, (ρpop = (e^2z^ -1)/(e^2z^ +1)). A Wilcoxon signed-rank test was applied to the Fisher-transformed correlation values to assess whether the population correlation differed significantly from zero.

### Acute slice recordings Animals

Acute slice recordings were performed in wild-type (C57BL/6J) mice of both sexes (postnatal days ∼50), housed 2–3 per cage on a 12 h light/dark cycle with ad libitum access to food and water. All procedures were carried out at the Institute of Experimental Medicine, Hungary (HUN-REN KOKI) and were approved by the institutional Ethical Committee for Animal Research (PE/EA/4118-4/2016) and conformed to Hungarian legislation (1998/XXVIII Law on Animal Welfare) and European Communities Council Directive 2010/63/EU on the protection of animals used for scientific purposes.

### Injections

To enable optogenetic activation of hippocampal principal cells, AAV9.CaMKII.hChR2(E123A).mCherry.WPRE.hGH (Addgene35506) was injected bilaterally into the dorsal hippocampus (AP: -1.4, -1.8, -2.3 mm; ML: 1.5, 2.0 mm from bregma, DV: 1.3 mm from dura; 100 nl virus per track). Red FluoSpheres (Thermo Fisher Scientific, diameter: 0.04 µm, excitation/emission: 580/605 nm) were injected into the MS (AP 0.9 mm, ML 0.0 mm from bregma, DV 3.5 and 3.8 mm from dura) to enable visually guided, targeted recordings from HS cells. In experiments involving the mCherry-expressing virus, red FluoSpheres were combined with green visualization of biocytin-filled cells. Despite their overlapping fluorescence, the FluoSpheres could be readily distinguished from the membrane-associated ChR2-mCherry signal based on their distinct appearance. In case of animals used for recording the ripple-related activity, green FluoSpheres were used with the same injection parameters as described above. As no additional fluorescent marker was present in these experiments, biocytin-filled cells could be visualized in red using streptavidin-Alexa Fluor 594.

### Acute slice preparation

Mice were deeply anesthetized with isoflurane and decapitated. Brains were rapidly removed into ice-cold, carbogenated (95% O2–5% CO2) cutting solution containing (in mM): 205 sucrose, 2.5 KCl, 26 NaHCO3, 0.5 CaCl2, 5 MgCl2, 1.25 NaH2PO4 and 10 glucose, for at least 30 min before recording. Coronal hippocampal slices (300 µm thick) were prepared using a vibratome (Leica VT1000S). Slices were then transferred to an interface-type holding chamber containing standard ACSF at 35°C that gradually cooled down to room temperature. The ACSF consisted of (in mM): 126 NaCl, 2.5 KCl, 26 NaHCO3, 2 CaCl2, 2 MgCl2, 1.25 NaH2PO4 and 10 glucose, continuously bubbled with carbogen. Cholinergic modulation of HS activity was induced by bath application of 5 μM carbachol.

### In vitro recording conditions

Recordings were performed under visual guidance using differential interference contrast (DIC) microscopy (Nikon FN-1) and a 40× water-immersion objective. After incubation for at least 2 h, slices were transferred individually into a submerged-type recording chamber perfused with oxygenated ACSF. Patch pipettes were pulled from thin-walled borosilicate glass capillaries with inner filament (OD 1.5 mm; Hilgenberg, Germany) using a PC-10 puller (Narishige, Japan). Signals were recorded with Multiclamp 700A or 700B amplifiers (Molecular Devices), digitized at 10–20 kHz via a USB-6353 DAQ board (National Instruments), and acquired with custom software written in C#.NET and VB.NET.

### In vitro cell identification

HS cells were targeted for recording based on somatic retrobead labeling (see Injections). For post-hoc identification, cells were filled with biocytin using an intracellular solution containing (in mM): 110 K-gluconate, 4 NaCl, 20 HEPES, 0.1 EGTA, 10 phosphocreatine, 2 ATP, 0.3 GTP, and 3 mg/ml biocytin (pH 7.3–7.35 adjusted with KOH; 285–295 mOsm/L). Following biocytin filling, slices were fixed in 4% paraformaldehyde in 0.1 M phosphate buffer (PB; pH 7.4) for ∼24 h and washed out several times in PB. Sections were then blocked with 10% normal goat serum in Tris-buffered saline (pH 7.4) and incubated with Alexa Fluor 488 or 594–conjugated streptavidin (1:1000; Invitrogen depending on the experimental condition (see Injections). Sections were mounted in Vectashield (Vector Laboratories), and recorded cells were identified by the presence of retrobeads in the soma.

### Optogenetic stimulation

For in vitro optogenetic stimulation, a 447 nm blue laser diode (Roithner LaserTechnik GmbH) coupled to a single optical fiber (Thorlabs) and a digital micromirror device (Polygon400, Mightex Systems, Toronto, Canada) integrated into the microscope optical path was used. Illumination patterns were defined as 50 × 50 pixel grids with an effective single-pixel size of ∼10 × 10 µm. Images with increasing pixel saturation were used to drive pyramidal cell activity and reveal the effect of progressively stronger excitatory synaptic input on HS cell firing (input–output transformation), and the same protocol was repeated in whole-cell mode to measure the synaptic charge corresponding to the level of spiking (measured in loose-patch configuration). Each saturation level (1, 5, 10, 20, 50, and 90%) was applied for several seconds (∼5 s, 2 ms single image duration), with active pixels randomly redistributed across the grid every 5 ms.

### Ripple recording in acute slices

After at least 1.5 h of incubation, 450 μm thick slices were transferred to a submerged recording chamber with dual ACSF superfusion above and below the tissue (3–3.5 ml/min per channel, 30–32°C), providing enhanced metabolic support^64^ enabling spontaneous sharp wave-ripple-like activity. LFP recordings were performed with ACSF-filled standard patch pipettes (3–6 MΩ) placed in str. pyramidale.

### Data analysis

All acute slice data were analyzed offline using custom-written programs in Delphi 6.0 (A.I.G.) and Matlab 8.5.0 and Python 2.7.0 (D.S.). All automatic detection steps were visually supervised. Spikes in loose-patch recordings were detected from traces high-pass filtered at 500 Hz using a threshold set to five times the standard deviation of the signal. For EPSC charge distribution, each baseline-subtracted EPSC was integrated in a 50 ms time window. For optogenetically evoked responses, cumulative EPSC charge was computed in 1 s windows. SWRs were detected in 30 Hz low-pass-filtered field recordings using a threshold of 4× the signal SD. For averaging and HS spike PSTHs, events were aligned to the negative peak of the ripple cycle nearest the SWR peak to preserve ripple phase.

### *In silico* model of the MS network with HS feedback

To examine how the HS feedback can shape MS neuronal activity and theta output, we extended a previously described conductance-based model of the MS pacemaker network^19^ with an additional GABAergic HS input. The model was implemented in the NEURON simulation environment^65^ using a fixed integration time step of 0.025 ms; the network parameters were generated and the simulation output was processed with custom MATLAB scripts.

Each MS neuron was represented as a single-compartment, fast-spiking cell (total membrane surface area 5000 µm², membrane capacitance 1 µF/cm²), based on a fast-spiking interneuron model^47^ and implemented as described previously^19^. The model contained a transient sodium current (Nas) generating action potentials, a delayed-rectifier potassium current (Kdr) mediating repolarization, a slowly inactivating D-type potassium current (Kd) shaping the after-hyperpolarization and the duration of bursts, an HCN-mediated hyperpolarization-activated current responsible for the characteristic sag response, and a passive leak current (leak conductance 0.1 mS/cm², leak reversal potential −60 mV). With this channel repertoire, model neurons reproduced both high-frequency tonic firing and theta/delta-range rhythmic bursting upon tonic excitation. The complete channel equations and parameters are provided in *Kocsis et al., 2022*^19^.

The network comprised twenty model neurons reciprocally connected by fast GABA-A synapses, implemented with the built-in NetCon and ExpSyn mechanisms of NEURON. A presynaptic spike (detected when the somatic membrane potential crossed a 0 mV threshold) evoked a step increase in synaptic conductance followed by an exponential decay, with a synaptic reversal potential of −70 mV. In the main, intra-MS-dominated configuration, neurons were interconnected at a 60% connection rate, obtained by randomly removing connections from a fully connected graph (self-connections excluded), with a mean synaptic delay of 7 ms, a mean decay time constant of 2 ms and a mean synaptic strength of 3 nS; a 10% coefficient of variation was applied independently to the delay, decay and weight of each synapse. Tonic excitation was delivered to every neuron as a somatic current step (IClamp). The baseline drive (60 pA, with 10% variance across neurons) was increased 1.4-fold (to 84 pA) to induce the theta state and subsequently returned to baseline; the onset of each current step was jittered across neurons to prevent artefactual synchronization by simultaneous stimulus onset, and a slow, delta-frequency (1 Hz) modulation of the tonic drive was superimposed during the non-theta periods.

The HS feedback was modeled as an additional GABA-A input that targeted a variable fraction of MS neurons (all neurons in the main configuration) and used the same synaptic mechanism and reversal potential as the intra-MS connections. The strength of the HS synapse was set relative to the intra-MS synaptic strength through a scaling factor, and the same 10% variance was applied to its weight, delay and decay. We simulated theta and non-theta states under three HS feedback regimes: a suppressed regime, in which the HS input was eliminated; a theta-rhythmic regime, in which the external GABAergic source delivered a theta-frequency rhythmic spike train (the baseline condition); and a non-rhythmic regime, in which the source discharged as a 50 Hz Poisson process.

Each simulation lasted 60 s and consisted of a 20 s baseline (non-theta) period, a 20 s period of elevated tonic excitation (induced theta) and a final 20 s return to baseline (non-theta). For each HS regime, the simulation was repeated 60 times, with all stochastic parameters (synaptic weights, delays and decay times, stimulation amplitudes, and the random assignment of intra-MS connections and HS targets) redrawn from the same distributions on every run. As a proxy for the septal output to the hippocampus, the spike trains of all neurons were convolved with a 50 ms Gaussian kernel and summed across the population. Theta and non-theta states of this population output were detected from the smoothed theta-to-delta power ratio (theta band 4–6 Hz, delta band 0.5–4 Hz), as described for the biological recordings previously^19^.

To compare the relative influence of intrinsic and feedback connectivity, in addition to the intra-MS-dominated configuration described above (60% intra-MS connectivity), we simulated a balanced configuration in which the summed intra-MS and HS synaptic strengths were comparable, obtained by reducing the intra-MS connection rate (to approximately 33%) and increasing the HS synaptic strength (to approximately 6.5-fold the intra-MS strength); the effect of HS elimination on the theta power of the network output was then compared between the two configurations.

### HS manipulation in anesthesia or freely behaving animals Animals

*In vivo* recordings (both anesthetized and freely behaving) were performed in adult Sst-IRES-Cre male mice, single housed after virus injection and kept on a 12 h light/dark cycle with ad libitum access to food and water. All procedures were carried out at the Institute of Experimental Medicine, Hungary (HUN-REN KOKI) and were approved by the institutional Ethical Committee for Animal Research and conformed to Hungarian legislation (1998/XXVIII Law on Animal Welfare) and European Communities Council Directive 2010/63/EU on the protection of animals used for scientific purposes.

### Virus injections

For selective manipulation of the HS pathway, we bilaterally injected the dorsal hippocampus of Sst-IRES-Cre mice with AAV2/5-Ef1a-DIO-ChR2-YFP-WPRE (2 x 2 x 150-200 nl) for optogenetic activation, or with AAV5-Ef1a-DIO-SwiChRca-TS-EYFP, (SwiChRca renamed later to SwiChR++), AAV-eNpHR-YFP, AAV-Arch-GFP, AAV5-ArchT-GFP or AAV-eArch3.0 (2 x 2 x 150-400 nl) for inhibition. Mice were anesthetized for virus injections with ketamine-xylazine mixture (i.p., 167 and 7 mg/kg, respectively). Following 3-20 weeks of the injection, animals were re-anesthetized for acute *in vivo* juxtacellular recordings (ChR2: n = 81; SwiChR: n = 41; Halo: n = 12; Arch: n = 5; eArch: n = 11; ArchT: n = 7 mice) or implanted with silicon probes for chronic recordings of the MS and hippocampal electric activity (ChR2: n = 10; SwiChR: n = 18 mice). A third cohort received bilateral AAV5-Ef1a-DIO-YFP injections (2 x 2 150 nl) and served as controls for optogenetic inhibition of the HS pathway in chronic recordings (n = 7 mice).

### Acute *in vivo* juxtacellular recordings

Mice were anesthetized by i.p. injection of 20% urethane (Sigma-Aldrich, St Louis, MO, USA; dose: 0.007 ml/g body weight). The anesthetic depth was verified by paw-withdrawal reflex. Body temperature was maintained with a homeothermic heating pad throughout the experiments. Mice were head-fixed in a stereotactic frame, the skull was exposed, and a cranial window was drilled to access the MS (coordinates: AP +0.9, ML+0.9, 14° angle) for juxtacellular recordings. Juxtacellular signals were recorded with filamented borosilicate micropipettes (20–45 MΩ) filled with 0.5 M NaCl containing 2% Neurobiotin (Vector Laboratories, Inc., Burlingame, CA, USA). After reaching the target zone (∼3000 μm below the brain surface), the recording pipette was advanced slowly (0.8–1 μm/s) using a micropositioner (EXFO, Quebec, Canada). The signal was band-pass filtered (0.1 - 5 kHz) and amplified (gain 500, 1000 or 2500) with an AxoClamp 2B amplifier (Axon Instruments, Foster City, CA, USA) and a LinearAmp signal conditioner (Supertech, Pécs, Hungary).

To enable optogenetic manipulation of HS fibers around the recorded cell, an optical fiber (105 μm diameter, 0.22 NA) was attached to the recording pipette. A second craniectomy was made above the dorsal CA1 (AP -2.5, ML -1.6 and 0° angle), and a monopolar stainless-steel wire was implanted 1200-1300 μm below the brain surface for local field potential (LFP) recordings. LFPs were filtered between 0.3 - 2 kHz or 5 kHz (gain 2000 or 5000) using an AC-coupled amplifier (BioAmp, Supertech). A silver wire placed under the neck skin served as common ground. All electrophysiological signals were digitized at 10 kHz using a Micro1401 mkII or a Micro3 1401 (Cambridge Electronic Design, Cambridge, UK) controlled by Spike2 software (Cambridge Electronic Design).

### Juxtacellular recording protocol

The MS single-cell activity and hippocampal LFP were recorded during control periods and during optogenetic manipulation of the HS pathway. Sensory-evoked hippocampal theta activity was elicited by tail pinch (for ∼30 s) using a crocodile clip with the serrated surface insulated by thin plastic tape. In ChR2-experiments, HS fibers in the MS were stimulated at 1, 5, 10, 20 and 50 Hz with blue (447 nm) light (5-8mW, Roithner Lasertechnik, Vienna, Austria). For 1 Hz stimulation, pulse width was 1, 5, 10, 20 or 30 ms; for 5–50 Hz stimulation, pulse width was 5 ms, except for one cell in which 10 and 20 Hz trains used 1 ms pulses, and three cells in which 10 or 20 Hz trains used 3 ms pulses. Trains at 1 Hz contained 60–120 pulses; 5 Hz trains contained 60–300 pulses (typically 100); 10 Hz trains contained 100–600 pulses (typically 200); 20 Hz trains contained 200–400 pulses (typically 200); and 50 Hz trains contained 250–500 pulses (typically 250), resulting in train durations mainly between 5–10 s.

In SwiChR experiments, blue (447 nm) illumination was applied for 5–10 s and followed, after a 0–7 s interval, by a 2–3 s yellow (593 nm; 8-10 mW, IkeCool Corporation, Los Angeles, USA) pulse. In experiments using Halo, Arch, eArch or ArchT, yellow light was delivered in single pulses lasting 10–60 s.

Stimulus trains (ChR2) or pulses (SwiChR, Halo, Arch, eArch and ArchT) were delivered 1-19 times, and were pseudorandomly interleaved within and outside tail pinch periods. After recording, the neurons were juxtacellularly labelled with Neurobiotin by applying 1-10 nA current (gradually increased until current modulation of neuronal firing was observed) through the recording pipette as 200 ms pulses (at 50% duty cycle), combined with a continuous cathodal DC current (0.5-0.9 nA).

### Histological verification following juxtacellular recordings

After successful labelling, animals were transcardially perfused with saline followed by 4% paraformaldehyde (survival time after labelling was 10-150 min). Brains were cut into 40–80 μm coronal sections on a vibratome (Leica Microsystems, Wetzlar, Germany). After extensive washing in 0.1 M phosphate buffer (pH 7.4), sections were transferred to Tris-buffered saline (TBS, Sigma-Aldrich, pH 7.4), and all subsequent washes and serum dilutions were carried out in TBS. Neurobiotin-filled neurons were visualized with CY3-conjugated streptavidin (Jackson ImmunoResearch, West Grove, PA, USA, catalogue no.: 016-160-084, dilution 1:1000). Parvalbumin-expression was assessed by immunofluorescence (primary: Rabbit anti Parvalbumin Lyophilized antiserum (Swant, Switzerland, catalogue no: PV27a, 1:1000); secondary: Dylight405-conjugated donkey anti-rabbit (Jackson ImmunoResearch, West Grove, PA, USA, catalogue no.: 711-475-152, dilution 1:400)). YFP in transduced cells was enhanced using anti-GFP immunofluorescence (GFP polyclonal antibody, Invitrogene, A10262, 1:1000 and Alexa488-conjugated goat anti chicken, Invitrogen, A11039, 1:1000).

In sections containing the hippocampus, additional staining for SST (primary: Polyclonal Guinea pig antiserum, lyophilized (Sysy, catalogue no.: 366004, 1:1000) and secondary: Cy3-conjugated donkey anti-guinea-pig (Jackson Immunoresearch, catalogue no.: 706-166-148, dilution 1:200)) was performed together with anti-YFP to quantify YFP-positive somata expressing SST (n = 6 ChR2 and n = 5 SwiChR mice). MS cells occasionally retrogradely labelled by the virus (injected to the hippocampus) were counted in n = 7 ChR2 animals and n = 14 SwiChR mice. The area (only in the CA1; only in the CA3; both in CA1 and CA3) of the hippocampal YFP-expression was determined in n = 39 ChR2 animals and n = 22 SwiChR animals. Sections were mounted on microscopic slides and covered with mounting medium (Vectashield, Vector Labs, Burlingame, CA, USA), fluorescence signals were examined with an Axioplan 2 microscope (Carl Zeiss, Oberkochen, Germany) equipped with a DP70 CCD-camera (Olympus, Tokyo, Japan) or with an A1R confocal laser scanning microscope (Nikon, Tokyo, Japan).

### Analysis of juxtacellular recordings

Spikes in the juxtacellular recordings were detected using Spike2 and all events were verified by visual inspection before export of spike times to Matlab for further analysis with custom-written scripts. Neuronal activity was quantified separately during hippocampal theta and non-theta states. Theta state was defined as tail pinch epochs; whereas non-theta was defined as periods outside the tail pinch epochs without dominant theta rhythm (periods with dominant theta outside tail pinches were excluded from the analysis). Dominant theta was identified when the theta score, defined as the (θ - δ)/ δ, exceeded the median value computed for the entire recording; θ and δ denote the spectral magnitudes (2.25-6 Hz and 0.9-2.25 Hz, resp.) obtained from L1-normalized wavelet-based decomposition of the hippocampal LFP after downsampling to 1 kHz, smoothing with a 10 ms overlapping moving average and z-scoring.

### Surgery for chronic recordings

Chronic silicon probe implantations were performed under isoflurane anesthesia, induced with ketamine-xylazine mixture (4:1; diluted 1:6 in Ringer’s lactate; i.p. 0.01 mg/g). Body temperature was maintained with a heating pad throughout the surgery. Mice were placed in a stereotaxic frame, the scalp was shaved, disinfected with Betadine, and infiltrated subcutaneously with lidocaine; eyes were protected with ophtalmic ointment (Laboratoires Théa, Clermont-Ferrand, France). After opening the skin and cleaning the skull, the head was leveled using bregma and lambda, and an adhesive bonding layer (OptiBond, Kerr, Brea, US) was applied to the cleaned bone to promote stable fixation of the implants. Before insertion, silicon probes were coated with red DiI (Thermo Fisher Scientific, Waltham, USA) to enable post-hoc histological reconstruction.

Craniectomies (∼1.5 mm diameter) were made above the MS (AP +0.9, ML +0.9, probe tilted 15° in coronal plane) and dorsal hippocampus (AP -3.1, ML -2.0, probe tilted 15° from dorsocaudal or AP -2.5 and ML -2 when implanted vertically) for probe placement. During implantation, the MS probe was lowered to just above the target (∼2.4 mm below the brain surface), and the hippocampal probe to 2.2 mm depth; both were mounted on custom-made microdrives for subsequent axial adjustment. In ChR2 mice, an additional craniectomy (AP -0.7, ML -1.7) was made to implant an optical fiber (105 μm core, 0.22 NA) targeting the fimbria at 1.9 mm depth (15° angle). In SwiChR animals with MS recordings, optical fibers (50 μm core, 0.22 NA) were attached to each shank of the MS silicon probe, with the fiber tips (etched by hydrofluoric acid to a pencil profile) positioned ∼75–100 μm above the uppermost recording site and secured with optical adhesive (Norland 61 or 81, Norland Products, Jamesburg, NJ, USA).

In YFP control mice (n = 3), silicon probes were implanted only into hippocampus (AP -2.5, ML +1.8, DV from brain surface 2 mm) and bilateral optical fibers (105 μm core, 0.22 NA) were implanted above the MS (coordinates AP +0.9, ML +-0.75, tilted 12°, DV from brain surface 3.0-3.2 mm). In n = 4 SwiChR mice and in n = 1 additional YFP control two hippocampal silicon probes were implanted (a multishank probe between AP -1.5 to -2.5 and ML +1 to ML +2.3, tilted and DV -1.6-1.7 mm from brain surface, and a single-shank probe in AP -3.0, ML +3.2 tilted 20°, DV -2mm) together with bilateral MS optical fibers. For control illumination, in n = 1 SwiChR animal with MS and hippocampal probes and in n = 2 SwiChR animals with hippocampal-only probes, additional optical fibers were attached to the head implant but terminating outside the skull (illumination denoted as “blind” in the figures), serving for control illuminations. The implanted optical fibers attenuated the laser light to 50-55% (checked before implantation). See Extended Data Table 2 for details of implanted silicon probes, optical fibers and chronic recordings used in this study.

Craniectomies were sealed with artificial dura (Cambridge NeuroTech Ltd, Cambridge, UK). Ground and reference screws were placed in the occipital bone and connected to the probes with insulated stainless-steel wires. The entire implant was shielded with copper mesh to reduce environmental electrical noise and encapsulated in dental acrylic (Paladur, Kulzer, Hanau, Germany). At the end of surgery, mice received buprenorphine (s.c., 0.045–0.1 μg/g) for analgesia, were monitored on a heating pad until full recovery, and were then returned to their home cages.

### Chronic recording procedures

Recordings were started after a one-week postoperative recovery period and habituation to handling and connectorization. MS probes were advanced in daily steps of 75-150 μm toward the target zone, defined by neurons with theta rhythm modulated firing; within the target zone the steps size was reduced to 45 μm. After each recording session, the MS probe was advanced further and recordings at the new depth were performed after a delay of at least 12-24 h.

If Buzsáki-type probes had been implanted in the hippocampus, their position was adjusted to record from the CA1 pyramidal layer, identified by the presence of ripples during immobility, complex spike patterns, and an increased multi-unit yield. For linear probes, the tip was placed below the pyramidal layer such that the recording sites spanned the pyramidal layer, no further adjustment was required. In YFP control mice (prioritized LFP recording over single-unit yield), the probes were left at their initial target depth.

Recordings were performed in the home cage to sample immobility and quiet wakefulness. Mice were then transferred to an unrewarded, 1 m x 0.09 m linear track with 0.04 m high walls to record activity during active exploration. In ChR2 mice, trains of 5ms blue light (447 nm, 15-22 mW) pulses at 5, 20 and 50 Hz were delivered to activate the HS pathway (repeated 4-23 times). In SwiChR and YFP mice, 10 s blue (447 nm, 15-22 mW) illumination was followed, after a 2 s interval, by a 1 s yellow (593 nm, 15-22 mW) pulse (repeated 2-28 times). In case of bilateral MS illumination, one side was illuminated by a 447 nm, the other side by a 473 nm blue laser, the yellow was fed only in the fiber with 447 nm illumination. Light delivery was triggered manually in pseudorandom order across the session.

Electrophysiological signals were recorded with a multiplexing data acquisition system (KJE-1001, Amplipex Ltd, Szeged, Hungary) at 20 kHz sampling rate. Animal position was tracked in 3D with a marker-based, high-speed (120 frames/s) motion capture system (Motive, OptiTrack, NaturalPoint Inc, Corvallis, OR, US). Spherical markers (3 mm OptiTrack facial markers) were attached to the corners of the head implant and to the headstage of the electrophysiology recording system.

### Histological verification

After completion of recordings, mice were perfused, and brains were processed as for the juxtacellular experiments. Silicon probe trajectories were visualized by red DiI fluorescence, the YFP expression was enhanced by anti-GFP immunostaining using the same protocol as for juxtacellular experiments. The sections were mounted and examined as in the juxtacellular experiments to confirm probe and optical fiber positions, and the transgene expression.

### Preprocessing for analysis of chronic recordings

Spike trains were detected and clustered with SpyKING Circus^66^ and manually curated in Phy2 in recordings with Buzsáki-type probes (MS and hippocampus when available). Only well-isolated units with clear refractory periods, consistent waveforms and well-separated amplitude parameter distributions on visual inspection were included in the analysis. Spike times were exported to Matlab and analyzed using the same pipeline as for juxtacellular spike trains. To minimize artefactual events from cluster merges or misdetections, events with instantaneous firing rates >800 Hz relative to the preceding spike were removed.

In hippocampal recordings, the channel with the largest ripple magnitude was selected as the pyramidal layer LFP; its anatomical location in dorsal CA1 was confirmed by histology. This pyramidal LFP served as the reference signal for the hippocampal state classification and spectral analyses. For ripple detection, data were analyzed at the original 20 kHz sampling rate, whereas all other LFP analyses used signals downsampled to 1 kHz.

Animal speed was derived from the 3D-tracked position. Markers attached to the implant and headstage defined a rigid body whose pivot was aligned to bregma in the Motive software; the resulting bregma coordinates were exported to Matlab, and speed was calculated as the Euclidean distance between successive frames divided by the frame interval (horizontal 2D speed for state definitions; 3D speed for speed modulated firing and LFP analyses). The speed trace was linearly interpolated to the 1 kHz LFP sampling rate and smoothed with an overlapping 0.5 s moving average for state definitions or 10-frame overlapping moving average for speed modulated firing and LFP. For state definitions, periods with missing tracking were filled by linear interpolation between the last and next valid samples. Movement thresholds were set to 4 cm/s in the linear track and 1 cm/s in the home cage. Short subthreshold segments (<0.2 s) were reclassified as movement, while suprathreshold segments shorter than 2 s were discarded.

In SwiChR mice, optical fibers were attached to each MS probe shank, and optogenetic illumination was delivered sequentially via individual fibers (only one fiber at a time; not all four fibers were necessarily used in every session). Only one manipulation site was retained for each neuron, chosen as the shank resulting in the lowest p value for firing rate differences between control and stimulation epochs in the linear track. The stimulation at the same shank was used for the analysis of the home cage part of the session.

Hippocampal states in chronic recordings were defined based on immobility/movement periods and the hippocampal dominant theta rhythm. For analyses in which stimulation and control epochs were pooled (for example, cross-correlogram calculations), we included only periods with movement in linear track recordings and periods with immobility and no dominant theta in home cage recordings. Dominant theta was defined as a theta (4–12 Hz) to delta (0.9–4 Hz) spectral magnitude ratio exceeded 1.5. For quantification of firing rate, bursting, phase coupling, and LFP spectral magnitudes, stimulation and control epochs were not pooled: each epoch was quantified separately, and for statistical comparisons, the mean across epochs was used for each neuron.

### Analysis pipeline

Spike trains from juxtacellular recordings and from units isolated with silicon probes were processed with a common analysis pipeline. Neurons were classified according to their theta-rhythmic firing based on autocorrelograms computed from the entire control recording period (applying state definition rules).

Stimulus epochs were defined as follows: in SwiChR experiments, as the duration of blue light pulses (excluding the subsequent yellow pulse and intervening gap); in Halo, Arch, eArch, and ArchT experiments, as the duration of the yellow light pulses; and in ChR2 experiments, as the interval from onset of the first pulse to offset of the last pulse within each stimulus train. Control epochs were defined as intervals of equal duration immediately preceding each stimulus epoch and were truncated if they overlapped with an earlier stimulus epoch.

Stimulus-induced changes in firing rate at the single-cell level were evaluated with paired t-tests across individual epochs. Firing rate was calculated as the total spike number in each epoch divided by the epoch duration. Based on the direction of significant firing rate changes in the theta state (linear track) for inhibitory opsins (SwiChR, Halo, Arch, eArch, ArchT) or in the non-theta (home cage) for ChR2 (20 Hz trains used for classification in anesthetized animals and 50 Hz trains in freely behaving animals), MS neurons were categorized as activated or suppressed. Neurons without significant firing-rate changes, or with fewer than 50 spikes in the entire recording, were classified as non-reactive. The effect size of the firing rate change was calculated as the difference in average firing rate between stimulation and control.

Theta modulation depth and burst parameters were calculated for each epoch, then averaged per neuron. Population data were compared using Wilcoxon signed-rank tests (theta versus non-theta, control versus stimulation) or two-way repeated-measures ANOVA (main effects of state and group, and state × group interaction), followed by Tukey’s honestly significant difference post hoc tests for multiple comparisons. Theta modulation depth was defined as the summed relative spectral magnitude in 5-12 Hz (in urethane experiments: 3-6 Hz) obtained by averaging of the L1-normalized wavelet-based decomposition of the autocorrelogram in 0 to +500 ms bins. Theta-bursts were defined as spike clusters containing at least 3 spikes with instantaneous firing rate over 30 Hz and lasting for maximum 0.15 s. Burst index was calculated as the ratio of spikes within the burst (including the first and the last spikes) to the total number of spikes. Burst occurrence was defined as the number of bursts divided by the epoch duration. Burst length was calculated as the interval between the first and the last spike in each burst, then averaged across bursts per epoch. Intra-burst frequency was calculated as the spike number per burst divided by burst length, then averaged per epoch.

The phase preference to hippocampal theta rhythm was analyzed only for neurons recorded in freely behaving animals. Stimulus train induced rhythmic firing and peristimulus time histograms (PSTHs) were examined in both urethane anesthetized and freely behaving ChR2 experiments. Ripple-aligned firing was assessed in home cage recordings, and speed modulated firing was examined in linear track. In animals with unreliable ripple detection (n = 3), phase preference was not analyzed. Parts of the freely behaving dataset have been reported previously^19,67^, but those studies did not include all the units analyzed here and their analysis focused partly on other parameters than those analyzed here.

### Rhythmicity categories

MS neurons were classified as non-rhythmic, theta-off, follower or pacemaker cells based on their theta-rhythmic firing during hippocampal non-theta and theta states. Clustering was performed separately on the two pooled datasets: juxtacellularly recorded neurons from SwiChR, Halo, Arch, eArch, ArchT and ChR2 experiments, and units from chronic recordings (SwiChR and ChR2 pooled). Rhythmicity categories were defined by combining clusters obtained from heuristic k-means clustering of autocorrelograms (ACGs), performed separately for non-theta (home cage) and theta (linear track) states using pooled control epochs and the corresponding state definitions, followed by visual inspection and merging of similar clusters.

Non-rhythmic cells lacked rhythmic ACG in both states; theta-off cells were rhythmic only in non-theta; followers were rhythmic only in theta; pacemakers exhibited rhythmic ACGs in both theta and non-theta. ACGs were computed in 1 ms bin size, smoothed with a 10 ms overlapping moving average and z-scored; ±200 ms was used for freely behaving data, and ±400 ms for juxtacellular recordings – accommodating to the slower theta under urethane anesthesia. When multiple files were available from the same neuron, the recording with the strongest theta-modulated ACG in the theta state was used. Neurons with ≤20 spikes in a given state were classified as non-firing in that state; neurons non-firing in both states were not assigned a rhythmicity category, whereas neurons lacking an ACG in only one state were classified based on the other state.

### Peristimulus time histograms

Peristimulus time histograms (PSTHs) were generated for ChR2 experiments using 1 ms bins aligned to the pulse onset (pulses in theta/linear track and non-theta/home cage were pooled across the recording). The firing probability was defined as the mean spike count per 1 ms bin divided by the total number of light pulses. Z-scored firing probability was used only for visualization, all quantitative analyses were performed on baseline corrected, non-normalized firing probabilities. Baseline correction was achieved by subtracting the mean firing probability in -50 to 0 ms preceding the pulse onset. The 0 to +200 ms interval was segmented to identify abrupt local changes (peaks or troughs). A negative peak was defined when the first detected peak was negative, its amplitude was defined as the minimum corrected firing probability within the corresponding segment, and its latency was the time of this minimum relative to ChR2 pulse onset. The peak duration was the number of bins within the negative-peak segment in which the corrected firing probability was below zero. The rebound was defined as the first positive peak occurring in any subsequent segment after the negative peak, its amplitude was defined as the maximum corrected firing probability within that segment, its latency was measured from the pulse onset. Net spike excess was calculated as the sum of corrected firing probability values above and below zero in the 0-200 ms window after pulse onset.

### Phase coupling to stimulation frequency

Phase coupling to optogenetic stimulation pulses was assessed in ChR2 experiments (5, 20 and 50 Hz). The pyramidal LFP was band-pass filtered around the stimulation frequency (4.5-5.5 Hz, 19-21 Hz and 49-51 Hz), and the phase of the Hilbert transform at each spike time was extracted to compute the circular mean phase and mean vector length. Non-uniformity of phase distributions was tested using Rayleigh test. For each neuron, the mean vector length was averaged across stimulation/control epochs, a circular mean of the mean phase was computed, and the evidence for phase locking across epochs was summarized using Fisher’s combined probability test on the Rayleigh p values.

### Phase coupling to hippocampal theta

Phase coupling of MS neurons to hippocampal theta was quantified in SwiChR and ChR2 experiments in freely behaving animals and – when available – also for hippocampal neurons in SwiChR experiments. Hippocampal neurons were not subclassified (pyramidal cells vs. interneurons), all well-isolated units with acceptable sorting quality were included. The pyramidal LFP was band pass filtered in 4-12 Hz, and the circular mean phase and mean vector length of the Hilbert-transform phase were computed at burst onset times only. The non-uniformity of phase distributions was tested using Rayleigh test. The parameters were averaged, summarized, and tested for phase locking using the same procedure as for phase locking to the stimulation frequency.

### Cross-correlograms

Cross-correlograms (CCGs) were computed in freely behaving experiments in which multiple MS neurons were recorded simultaneously. CCGs were calculated using 1 ms bins, bin counts were normalized by the number of trigger spikes to get firing probability and then smoothed with a 10 ms overlapping moving average. Spikes across control/stimulation epochs (applying relevant movement and dominant theta criteria, see state definitions above) were pooled for the calculation. Z-scored CCGs (±200 ms window) from control linear track data were grouped by k-means clustering, followed by visual merging of similar clusters. For each neuron pair, a single CCG was used in clustering and subsequent analyses, selected based on the highest theta comodulation index in the linear track data.

Clusters were classified as rhythmically or non-rhythmically coupled based on their population-level theta comodulation index. The theta comodulation index was defined as the summed relative spectral magnitude in 5–12 Hz, obtained from the L1-normalized wavelet decomposition of the z-scored CCG in the 0 to +500 ms range. The coupling index was calculated as the difference between the central amplitude (mean firing probability over ±10 ms lag) and the baseline (mean firing probability in the −3 to −2 s and +2 to +3 s lag) of the non–z-scored CCG. The asymmetry index was defined as the difference between the mean amplitude in the +1 to +200 ms and in the −200 to 0 ms lags (for statistical comparisons, its absolute value was used). Relative changes in theta comodulation were expressed as (stimulation − control)/(stimulation + control). The CCG phase was obtained as the Hilbert-transform phase of the 5–12 Hz–filtered, z-scored CCG at zero lag. Cluster-level differences in theta comodulation, coupling and asymmetry indices (in linear track and home cage) were tested with two-way repeated measures ANOVA (group x state) followed by Tukey’s honestly significant difference post hoc tests for multiple comparisons.

CCGs were excluded if the baseline firing probability was below 0.0005 or if no corresponding stimulation-epoch CCG was available. Additional cluster-specific exclusion criteria were applied to remove weakly coupled pairs (non-rhythmic cluster with negative coupling index: central amplitude >= −0.9; non-rhythmic cluster with positive coupling index: central amplitude <= 2.5; asymmetric clusters: absolute asymmetry index <= 0.05). All excluded CCGs were reassigned to cluster #8, which contained weakly coupled pairs. This cluster, together with the rhythmic cluster #3 with weak theta comodulation were not used in further analysis. The remaining CCG pairs were compared between control and stimulation epochs using Wilcoxon signed-rank tests.

### Ripple detection

Ripples were detected in home cage recordings of SwiChR and YFP mice (in ChR2 animals, ripple detection was used only to identify the pyramidal layer; no further ripple analyses were performed because the ChR2 stimulation rhythm strongly entrained hippocampal LFP). In n = 3 SwiChR mice, ripple detection was unreliable, and these animals were excluded from ripple-related analyses.

Ripples were defined as events in which the envelope of the 150–250 Hz–filtered LFP (Hilbert magnitude) exceeded the mean by >2 SD, with a peak >4 SD above the mean. Peaks had to be separated by at least 15 ms; segments whose end and subsequent start were <10 ms apart were merged, keeping a single peak (the one with larger amplitude). Ripple duration (time between first and last sample above 2 SD) had to be 20-150 ms, and ripple frequency (dominant frequency >100 Hz from Welch’s power spectral density of the ripple segment) had to fall between 150-250 Hz. Ripple rate was defined as the number of ripple events during the pooled control/stimulation epochs divided by the epoch duration.

### Quantification of ripple modulated firing

Spikes of MS and hippocampal neurons were aligned to ripple peaks and counted in 5 ms bins. For MS neurons, spike trains only with SwiChR mediated manipulation at one selected optical fiber were used as in other parts of the analysis; for hippocampal neurons, all home cage stimulation sessions were pooled. For each neuron, the spike-count histogram aligned to ripple peaks was first smoothed with a 10-bin overlapping moving average and then averaged across ripple events to get firing probability trace. This trace was baseline-corrected by subtracting the mean firing probability in the −500 to −400 ms window before the ripple peak. To quantify ripple modulated firing, baseline-corrected firing probabilities were summed over the ±50 ms window. Ripple events occurring in the control/stimulation epochs were pooled across epochs for quantification of changes upon the optogenetic manipulation.

### Detection of ripple modulated neurons

Neurons negatively modulated by ripples were identified using Ward’s agglomerative clustering on two features: the timing of negative peaks relative to ripple onset and their durations. Data from both MS and hippocampal neurons were included in the clustering; firing around ripple onsets in entire control recording (not only stimulus-preceding control epochs) were used. Negative peaks were defined as local minima exceeding at least the half of the minimum of the z-scored, 10-bin-smoothed firing probability around ripple onset; peak duration was measured between the points where the signal crossed a threshold at the half peak prominence. The cluster with neuronal population negatively modulated by ripples was selected by visual inspection, the cluster was refined by excluding neurons with mean central (0-50 ms after the ripple start) z-scored firing probability >0. Further, in evaluation of optogenetic stimulation effects, only neurons with central z-scored firing probability <- 0.5 when calculated from pooled stimulus-preceding control epochs were used.

Positively modulated neurons were identified with the same procedure, but using the timing and duration of positive peaks that exceeded at least the half of the maximum of the z-scored, smoothed firing probability. The corresponding cluster was refined by excluding neurons with central z-scored firing probability ≤0 and neurons whose positive peak fell outside the -100 to +250 ms window around ripple onset. In evaluation of the optogenetic stimulation effects, only neurons with central z-scored firing probability >0.5 within the pooled stimulus-preceding control epochs were used.

### Speed modulated firing

To assess speed modulation of MS firing, we used individual theta cycles of the pyramidal LFP as temporal reference frames. Theta cycles were defined from the phase of the Hilbert transform of the 5–12 Hz–filtered pyramidal LFP between −180 and +180 degrees, and only non-abrupt cycles (crossing the 0 rad, detected as |phase| < 0.1 rad) with durations between 20 and 250 ms were included. For each neuron, only cycles containing at least one spike were analyzed. Within each cycle, we computed the mean firing rate and the animal’s mean 3D speed. Spearman’s correlations between firing rate and speed were calculated separately for pooled stimulation/control epochs. As a control analysis, Spearman’s correlations were also computed per individual stimulation/control epochs (without pooling).

For each neuron, the epoch-wise correlation coefficients were combined by Fisher’s z-transform to get a single ρ estimate, and the associated p value was computed using Fisher’s combined probability test. Neurons with p < 0.05 in both the pooled and epoch-wise analyses during control periods were classified as speed-ON (ρ > 0) or speed-OFF (ρ < 0). Neurons that were not significantly speed-modulated during control, but showed significant correlations in both pooled and epoch-wise analyses during stimulation, were assigned to a third category, speed-STIM.

The fraction of speed-ON, speed-OFF, and speed-STIM neurons were compared across rhythmicity classes (non-rhythmic, theta-off, follower, pacemaker) using chi-square tests followed by post hoc Fisher’s exact tests.

### LFP entrainment by ChR2

To assess rhythmic entrainment of LFP oscillations by ChR2 stimulation, we computed stimulus-triggered LFP averages by averaging z-scored LFP segments aligned to individual ChR2 pulses. For each pulse, the L1-normalized wavelet decomposition of the LFP was obtained, and the average spectral magnitude and phase across pulses were calculated in peripulse windows (±1 s); circular means were used for phase, and spectral phase variance was quantified as circular variance averaged across pulses.

### Theta score

Theta (5-12Hz in freely moving animals, 2.25-6 Hz under urethane anesthesia) and delta (0.9-3Hz in freely moving animals, 0.9-2.25 Hz under urethane anesthesia) band magnitudes were computed from the L1-normalized wavelet decomposition of the LFP as the mean spectral magnitude. The theta score was defined as (theta magnitude−delta magnitude)/delta magnitude, and the effect size of stimulation was calculated as the difference in theta score between stimulation and control epochs. Theta score (and theta, delta band magnitudes) were calculated by individual control/stimulation epochs, then averaged per recording.

### Speed modulated theta score

The average 3D speed of the animal was computed on a theta cycle–by–theta cycle basis, as described for the analysis of speed modulated firing. Theta cycles were curated using the same criteria, except that the presence of spikes was not required here. For each cycle, the average theta score was obtained. Speed values were binned in 0.5 cm/s steps, and the relationship between binned speed (restricted to 5–10 cm/s) and the corresponding theta score during control versus stimulation was evaluated with a three-way repeated-measures ANOVA (speed x theta score with control vs. stim. as rm factor). Theta cycle duration was analyzed in the same way using a three-way repeated-measures ANOVA (speed x theta cycle duration with control vs. stim. as rm factor). SwiChR animals excluded from ripple analysis (n = 3) were also omitted from the analysis of speed modulated theta score.

### Cross-frequency coupling

To detect changes in theta-coupled higher-frequency oscillations, we quantified cross-frequency coupling using the modulation index between theta phase and high-frequency amplitude, as described by Tort et al.^68^ Theta phase was extracted from the Hilbert transform of the 5-12 Hz filtered pyramidal LFP and binned in 2π/50 steps; the amplitude of 15-200 Hz oscillations was derived from the L1-normalized wavelet-based decomposition. The modulation index was averaged within slow gamma (40-60 Hz), mid-gamma (60-90 Hz) and fast gamma (120-150 Hz) bands individually for each control/stimulation epoch, then finally averaged across epochs for each recording.

## Notes

### Competing Interest Statement

The authors have declared no competing interest.

