## Supplemental figures and legends for "Hippocampo-septal inhibitory feedback controls the medial septal circuit as a function of brain state, pyramidal cell activity and movement speed"

**EXTENDED DATA – FIGURE LEGENDS**

**Ext. Data Figure 1. Cholinergic modulation of HS neurons**

**a,** Example loose-patch recording from an identified HS cell (same targeting strategy as in Fig. 1f) during baseline and after increasing the cholinergic tone by bath application of 5 µM carbachol (CCh).

**b,** Firing rate change of HS cells in response to CCh (n=3 cells from 3 animals).

**c-d,** Current injection protocol of an identified HS cell during baseline and under CCh (c), and in the presence of fast excitatory synaptic blockers (d; 50 µM AP5 and 20 µM NBQX). Spike rasters are shown below. Note the CCh-induced depolarization of the resting membrane potential, and the barrage of excitatory synaptic potentials, which are strongly reduced by excitatory blockers, indicating that elevated cholinergic tone drives HS cell firing indirectly via pyramidal cell activation.

**e,** Resting membrane potential of identified HS cells in response to CCh, with and without fast excitatory blockers (n=6/6 cells from 6 animals). CCh-evoked depolarization was abolished in the presence of glutamatergic blockade.

**Ext. Data Figure 2. Histological verification of transduction of SST neurons**

**a,** Transduction of hippocampal neurons in Sst-Cre mice. Anti-YFP (left), anti-SST (middle) and merged (right) immunofluorescence images from dorsal CA1 region. a1, A SST-negative transduced principal neuron; a2-3, SST-positive neurons (a2, transduced; a3, unlabeled) in the stratum oriens.

**b,** Median number of transduced somata per animal (YFP-positive-only, SST-positive-only, and double positive cells) in CA1, CA3 and dentate gyrus (GD); top, ChR2 mice (n = 6); bottom, SwiChR mice (n = 5).

**c-d,** Fraction of YFP-only, SST-only and double positive neurons across (c) and within (d) hippocampal layers in CA1, CA3, and GD (ori, str. oriens; pyr, pyramidale; rad, radiatum; lam, lacunosum-moleculare; luc, lucidum; mol, moleculare; gra, granulosum; hil, hilus). Top, ChR2 mice; bottom, SwiChR mice.

**e,** Example hippocampal images illustrating CA1-restricted versus CA1-CA3-DG wide transduction.

**f,** Proportion of animals with CA1-only, CA3-only, or combined CA1+CA3 transduction (right, ChR2, n = 25; left, SwiChR, n = 11).

**g,** Effect size (change in firing rate, left; change in burst index, right) in CA1-only, CA3-only and CA1+CA3 transduced animals during non-theta and theta states under urethane anesthesia. One-way ANOVA, group (CA1-CA3) effect: firing rate, ChR2 20Hz, non-theta $F(2,24) = 1.5004$, $p = 0.2432$; theta $F(2,22) = 1.8593$, $p = 0.1794$; SwiChR, non-theta $F(2,19) = 0.7559$, $p = 0.4832$; theta $F(2,18) = 0.8553$, $p = 0.4417$; burst index, ChR2 20Hz, non-theta $F(2,22) = 0.1886$, $p = 0.8294$; theta $F(2,20) = 0.5884$, $p = 0.5645$; SwiChR, non-theta $F(2,19) = 1.3697$, $p = 0.2782$; theta $F(2,17) = 1.6093$, $p = 0.2290$; one outlier in the SwiChR non-theta, CA1+CA3 firing rate group is not shown.

**h,** Left, overview of the MS showing HS axons after hippocampal injection; right, higher-magnification view of a retrogradely labelled neuron in the horizontal limb of the diagonal band (HDB) (corresponding to the white boxed region in the left panel).

**i,** Top, number of labelled MS neurons per mouse after hippocampal injections (left, ChR2, n = 7; right, SwiChR, n = 14). Bottom, MS neuron counts as a function of days after hippocampal injection (Spearman's correlation: ChR2 $\rho = -0.23424$, $p = 0.62302$; SwiChR $\rho = 0.54621$, $p = 0.043302$).

**j,** Relative firing rate change ((stim - ctrl)/(stim + ctrl)) from linear track recordings of freely behaving animals plotted against days after injection. Spearman's correlation for |effect|: ChR2 $\rho$ = -0.22057, p = 0.0015652; SwiChR $\rho$ = 0.010941, p = 0.82164.

**Ext. Data Figure 3. Rhythmicity groups of MS neurons**

**a-b,** Autocorrelogram (ACG) clusters of MS neurons under urethane anesthesia during hippocampal non-theta (a; NTH1-4) and theta (b; TH1-4) states for all juxtacellularly recorded cells (including ChR2, SwiChR, Halo, Arch, eArch and ArchT experiments).

**c,** Color-coded cell counts by the combinations of non-theta and theta ACG clusters; boxed combinations define rhythmicity groups. Cells lacking sufficient spikes to generate ACGs in both states were excluded. Non-rhythmic cells were defined as cells not classified as pacemaker, follower or theta-off.

**d,** ACGs grouped by rhythmicity class (non-rhythmic, theta-off, follower, pacemaker) during non-theta (top) and theta (bottom).

**e-f,** ACG clusters of MS neurons in free behavior during home cage (e; HC1-4) and linear track (f; LT1-4) recordings including ChR2 and SwiChR experiments.

**g,** Color-coded neuron counts by the combinations of home cage and linear track ACG clusters; boxed combinations define rhythmicity groups. Cells lacking sufficient spikes to generate ACGs in both states were excluded. Non-rhythmic cells were defined as cells not classified as pacemaker, follower or theta-off.

**h,** ACGs grouped by rhythmicity class (non-rhythmic, theta-off, follower, pacemaker) during home cage (top) and linear track (bottom); color bar applies for a,b,d and e,f,h.

**Ext. Data Figure 4. Response to ChR2 stimulus pulses**

**a-e,** Amplitude (a), latency (b) and duration (c) of the first negative PSTH peak, rebound amplitude (d) and latency (e) by cell group and hippocampal state under anesthesia. Right-hand boxes summarize two-way RM ANOVA tests for rhythmicity-group effects (N, non-rhythmic; O, theta-off; F, follower; P, pacemaker), asterisks above box plots indicate significant differences between states (theta/non-theta) within the group. Two-way RM ANOVA tests: a, state $F(1,26) = 0.0184$, $p = 0.8931$; group x state $F(3,26) = 1.3941$, $p = 0.2669$; group $F(3,26) = 3.2546$, $p = 0.0377$; b, state $F(1,26) = 0.0043$, $p = 0.9481$; group x state $F(3,26) = 1.1526$, $p = 0.3466$; group $F(3,26) = 2.5001$, $p = 0.0817$; c, state $F(1,26) = 5.1072$, $p = 0.0334$; group x state $F(3,26) = 0.5238$, $p = 0.6698$; group $F(3,26) = 1.6584$, $p = 0.2004$; d, state $F(1,16) = 1.811$, $p = 0.1971$; group x state $F(3,16) = 0.9005$, $p = 0.4626$; group $F(3,16) = 1.0359$, $p = 0.4033$; e, state $F(1,16) = 7.5752$, $p = 0.0142$; group x state $F(3,16) = 1.0692$, $p = 0.3899$; group $F(3,16) = 0.9455$, $p = 0.442$.

**f-j,** Amplitude (a), latency (b) and duration (c) of the first negative PSTH peak, rebound amplitude (d) and latency (e) by cell group and hippocampal state in free behavior. Right-hand boxes summarize two-way RM ANOVA tests for rhythmicity-group effects (N, non-rhythmic; O, theta-off; F, follower; P, pacemaker). Two-way RM ANOVA tests: a, state $F(1,139) = 3.9958$, $p = 0.0476$; group x state $F(3,139) = 5.65$, $p = 0.0011$; group $F(3,139) = 3.4471$, $p = 0.0185$; b, state $F(1,139) = 7.5605$, $p = 0.0068$; group x state $F(3,139) = 1.1983$, $p = 0.3128$; group $F(3,139) = 5.1497$, $p = 0.0021$; c, state $F(1,139) = 13.9049$, p

= 2.7914x10-4; group x state $F(3,139) = 0.2559$, $p = 0.8570$; group $F(3,139) = 4.3674$, $p = 0.0057$; d, state $F(1,94) = 3.1318$, $p = 0.08$; group x state $F(3,94) = 2.5776$, $p = 0.0583$; group $F(3,94) = 4.575$, $p = 0.0049$; e, state $F(1,94) = 1.8273$, $p = 0.1797$; group x state $F(3,94) = 1.7184$, $p = 0.1685$; group $F(3,94) = 0.6669$, $p = 0.5744$.

**k,** PSTHs (5 Hz) grouped by response class defined from ChR2 20 Hz stimulation in non-theta.

**l,** Net spike excess by response group (S, suppressed, A, activated, N, unaffected) and state (non-theta and theta). Two-way RM ANOVA: state $F(1,33) = 2.7832$, $p = 0.1047$; group x state $F(2,33) = 8.3229$, $p = 0.0012$; group $F(2,33) = 7.3766$, $p = 0.0022$.

**m,** PSTHs (5 Hz) grouped by response class defined from ChR2 50 Hz stimulation in home cage.

**n,** Net spike excess by response group and state. Two-way RM ANOVA: state $F(1,200) = 0.3077$, $p = 0.5757$; group x state: $F(2,200) = 0.9963$, $p = 0.3711$; group: $F(2,200) = 4.2709$, $p = 0.0153$

**o,** Control firing rate (preceding the stimulus train) and baseline firing rate in the PSTH (0-50 ms before pulse onset) of suppressed neurons. Filled circles mark neurons with net spike excess > 0, despite exhibiting overall suppression during the full stimulus train (in 50 Hz stimulation).

**p**, Fraction of activated and suppressed MS cells across increasing stimulation frequencies of ChR2 pulses during hippocampal non-theta (urethane anesthesia, left) and in home cage (free behavior, right).

**Ext. Data Figure 5. Bursting changes of MS neurons upon optogenetic manipulations**

**a-b**, Burst index of MS neurons (grouped by rhythmicity and reactivity; the activated neurons during ChR2 stimulation are not shown due to low sample size) during ChR2 50 Hz (a) and SwiChR (b) in free behavior; Wilcoxon signed-rank tests (control vs stimulation): a, ChR2 50 Hz, HC, suppressed, N, $p = 2.0007 \times 10^{-11}$, O, $p = 0.008$, F, $p = 8.12 \times 10^{-7}$, P, $p = 0.0002$; non-reactive, N, $p = 0.045$, O, $p = 0.25$, F, $p = 0.625$, P, $p = 1$; LT, suppressed, N, $p = 6.24 \times 10^{-7}$, O, $p = 0.016$, F, $p = 4.51 \times 10^{-6}$, P, $p = 0.0004$; non-reactive, N, $p = 4.62 \times 10^{-5}$, O, $p = 0.5$, F, $p = 0.5625$, P, $p = 1$; b, SwiChR, HC, activated, N, $p = 0.765$, O, $p = 0.5$, F, $p = 0.297$, P, $p = 0.252$; suppressed, N, $p = 0.407$, O, $p = 0.031$, F, $p = 0.266$, P, $p = 0.021$; non-reactive, N, $p = 0.908$, O, $p = 0.049$, F, $p = 0.87$, P, $p = 0.038$; LT, activated, N, $p = 0.04$, O, $p = 0.5$, F, $p = 0.016$, P, $p = 0.001$; suppressed, N, $p = 0.004$, O, $p = 0.063$, F, $p = 0.00098$, P, $p = 0.00098$; non-reactive, N, $p = 0.557$, O, $p = 0.97$, F, $p = 0.334$, P, $p = 0.139$.

**c-d,** Burst occurrence; Wilcoxon signed-rank tests: c, ChR2 50 Hz, HC, suppressed, N, $p = 2.04 \times 10^{-12}$, O, $p = 0.004$, F, $p = .65 \times 10^{-7}$, P, $p = 0.0002$; non-reactive, N, $p = 0.01$, O, $p = 0.25$, F, $p = 0.625$, P, $p = 1$; LT, suppressed, N, $p = 9.03 \times 10^{-6}$, O, $p = 0.023$, F, $p = 1.35 \times 10^{-6}$, P, $p = 0.0003$; non-reactive, N, $p = 0.018$, O, $p = 0.5$, F, $p = 1$, P, $p = 1$; d, SwiChR, HC, activated, N, $p = 0.765$, O, $p = 0.5$, F, $p = 0.109$, P, $p = 0.277$; suppressed, N, $p = 0.021$, O, $p = 0.047$, F, $p = 0.38$, P, $p = 0.034$; non-reactive, N, $p = 0.828$, O, $p = 0.088$, F, $p = 0.318$, P, $p = 0.088$; LT, activated, N, $p = 0.0002$, O, $p = 0.5$, F, $p = 0.016$, P, $p = 0.0001$; suppressed, N, $p = 3.53 \times 10^{-5}$, O, $p = 0.031$, F, $p = 0.00098$, P, $p = 0.00049$; non-reactive, N, $p = 0.372$, O, $p = 0.733$, F, $p = 0.214$, P, $p = 0.256$.

**e-f,** Intra-burst frequency; Wilcoxon signed-rank tests: e, ChR2 50 Hz, HC, suppressed, N, $p = 0.858$, O, $p = 0.945$, F, $p = 0.358$, P, $p = 0.004$; non-reactive, N, $p = 0.637$, O, $p = 0.75$, F, $p = 0.8125$; LT, suppressed, N, $p = 0.337$, O, $p = 0.195$, F, $p = 0.122$, P, $p = 0.004$; non-reactive, N, $p = 0.718$, O, $p = 0.25$, F, $p = 0.156$, P, $p = 1$; f, SwiChR, HC, activated, N, $p = 0.422$, O, $p = 1$, F, $p = 0.375$, P, $p = 0.252$; suppressed, N, $p = 0.082$, O, $p = 0.625$, F, $p = 0.47$, P, $p = 0.557$; non-reactive, N, $p = 0.927$, O, $p = 0.011$, F, $p = 0.455$, P, $p = 0.359$; LT, activated, N, $p = 0.339$, O, $p = 0.5$, F, $p = 0.078$, P, $p = 0.002$; suppressed, N, $p = 0.295$, O, $p = 0.188$, F, $p = 0.519$, P, $p = 0.003$; non-reactive, N, $p = 0.759$, O, $p = 1$, F, $p = 0.425$, P, $p = 0.18$.

**g-h,** Burst length; Wilcoxon signed-rank tests: g, ChR2 50 Hz, HC, suppressed, N, $p = 5.34\times10^{-5}$, O, $p = 0.547$, F, $p = 0.001$, P, $p = 0.011$; non-reactive, N, $p = 0.451$, O, $p = 0.5$, F, $p = 0.188$; LT, suppressed, N, $p = 8.44\times10^{-7}$, O, $p = 0.742$, F, $p = 0.0096$, P, $p = 0.586$; non-reactive, N, $p = 0.0078$, O, $p = 0.25$, F, $p = 0.563$, P, $p = 1$; h, SwiChR, HC, activated, N, $p = 0.038$, O, $p = 0.5$, F, $p = 0.813$, P, $p = 0.208$; suppressed, N, $p = 0.465$, O, $p = 0.813$, F, $p = 0.129$, P, $p = 0.084$; non-reactive, N, $p = 0.844$, O, $p = 0.356$, F, $p = 0.132$, P, $p = 0.252$; LT, activated, N, $p = 0.028$, O, $p = 1$, F, $p = 0.0156$, P, $p = 0.391$; suppressed, N, $p = 0.0099$, O, $p = 0.4375$, F, $p = 0.003$, P, $p = 0.424$; non-reactive, N, $p = 0.998$, O, $p = 0.734$, F, $p = 0.618$, P, $p = 0.106$.

**Ext. Data Figure 6. CCG clusters**

**a,** Cross-correlogram (CCG) clusters obtained by k-means clustering (with visual curation) from linear track recordings; corresponding home-cage CCGs for the same neuron pairs are shown in rows, with white spaces indicating missing home-cage data due to low spike counts. Each neuron pair was represented only once, chosen by the highest control theta comodulation index (in the linear track). Minimal feature criteria (see b,c) were applied to exclude weakly coupled pairs from the clusters; pairs failing these criteria are grouped in cluster #8 (mixed pairs with minimal detectable modulation).

**b,** Schematic illustration of the key CCG features.

**c,** Theta comodulation index, coupling index (separately for negative and positive peaks) and asymmetry index for each CCG cluster. For each metric, exclusion thresholds were applied so that pairs with minimal feature values were omitted from statistical comparisons (see Methods for details). Left, box plots of CCG features by clusters (linear track and home cage); green shading highlights outstanding clusters regarding the given CCG feature; * above box plots marks p-values from the Wilcoxon signed-rank tests. Note that cluster #3 contains weakly theta modulated CCGs and was therefore not included in the opsin-effect analysis (Fig. 5). Right, summary of post hoc comparisons following the RM ANOVA for groups. Asterisks above the box plots indicate significant state difference (linear track vs home cage) within the given cluster. Statistics: theta comodulation index, RM ANOVA: state $F(1,2983) = 847.34$, $p = 3.4526\times10^{-164}$, group x state $F(7,2983) = 269.02$, $p = 2.4987\times10^{-311}$, group $F(7,2983) = 358.35$, $p = 0$; negative coupling index, RM ANOVA: state $F(1,5805) = 29.775$, $p = 5.0524\times10^{-8}$, group x state $F(6,5805) = 18.972$, $p = 5.3539\times10^{-22}$, group $F(6,5805) = 21.759$, $p = 1.9394\times10^{-25}$; positive coupling index, RM ANOVA: state $F(1,5294) = 2.021$, $p = 0.1552$, group x state $F(4,5294) = 44.803$, $p = 4.6979\times10^{-37}$, group $F(4,5294) = 212.02$, $p = 5.6251\times10^{-169}$; asymmetry index, RM ANOVA: state $F(1,5195) = 0.5018$, $p = 0.4787$, group x state $F(7,5195) = 19.433$, $p = 7.7633\times10^{-26}$, group $F(7,5195) = 18.5929$, $p = 1.2276\times10^{-24}$.

**d,** Composition of each CCG cluster by rhythmicity group (N, non-rhythmic, O, theta-off, F, follower, P, pacemaker, 0, uncategorized due to insufficient spikes for ACG clustering). The expected distribution corresponds to the overall rhythmicity distribution of individual neurons prior to partitioning into CCG clusters. Cluster #8 is not shown.

**Ext. Data Figure 7. Hippocampal effects under anesthesia and free behavior**

**a,** Theta (2.25-6 Hz, top) and delta (0.9-2.25 Hz, bottom) magnitudes of hippocampal LFP under urethane anesthesia. Theta magnitude, Wilcoxon signed-rank tests: SwiChR non-theta, $p = 0.30834$; SwiChR theta, $p = 0.0098322$; ChR2 5 Hz non-theta, $p = 0.23066$; ChR2 5 Hz theta, $p = 0.00012094$; ChR2 20 Hz non-theta, $p = 0.0001906$; ChR2 20 Hz theta, $p = 0.41096$; ChR2 50 Hz non-theta, $p = 8.1854\times10^{-5}$; ChR2 50 Hz theta, $p = 0.2865$. Delta magnitude: SwiChR non-theta, $p = 0.5119$; SwiChR theta, $p = 0.044482$; ChR2 5 Hz non-theta, $p = 0.63623$; ChR2 5 Hz theta, $p = 2.2305\times10^{-6}$; ChR2 20 Hz non-theta, $p = 0.53801$; ChR2 20 Hz theta, $p = 0.41096$; ChR2 50 Hz non-theta, $p = 0.026801$; ChR2 50 Hz theta, $p = 0.91067$.

**b,** Theta score during ChR2 stimulation at 5, 20 and 50 Hz under urethane anesthesia; Wilcoxon signed-rank tests: 5Hz non-theta, $p = 0.043902$; 5Hz theta, $p = 1.3058\times10^{-6}$; 20Hz non-theta, $p = 0.041284$; 20Hz theta, $p = 0.36139$; 50Hz non-theta, $p = 0.018694$; 50Hz theta, $p = 0.38971$.

**c**, Theta score during HS blockade under urethane anesthesia; Wilcoxon signed-rank tests: non-theta, $p = 0.43452$; theta, $p = 0.26425$.

**d,** Effect size of stimulations on the theta score (stim-ctrl) in urethane anesthesia; error bars indicate mean ±SEM.

**e,** Theta (5-12 Hz, top) and delta (0.9-3 Hz, bottom) magnitudes of hippocampal LFP. Theta magnitude, two-way RM ANOVA (SwiChR vs YFP, stimulation effect): home cage (HC), $F(1,136) = 2.8712$, $p = 0.0925$; linear track (LT): $F(1,140) = 6.2728$, $p = 0.0134$; multiple comparison(control vs stimulation) within group, HC SwiChR, $p = 0.0023$, HC YFP, $p = 0.4059$; LT SwiChR, $p = 1.0615\times10^{-10}$, LT YFP, $p = 0.9713$; Wilcoxon signed-rank for ChR2: 5Hz HC, $p = 0.026611$; 5Hz LT, $p = 0.003418$; 20Hz HC, $p = 0.00024414$; 20Hz LT, $p = 0.00024414$; 50Hz HC, $p = 0.00073242$; 50Hz LT, $p = 0.057373$. Delta magnitude, two-way RM ANOVA (SwiChR vs YFP), HC: $F(1,136) = 0.0543$, $p = 0.8161$; LT: $F(1,140) = 12.3047$, $p = 6.0779\times10^{-4}$; multiple comparison (control vs stimulation) within group, HC SwiChR, $p = 1.885\times10^{-5}$, HC YFP, $p = 0.2818$; LT SwiChR, $p = 2.8339\times10^{-5}$, LT YFP, $p = 0.0256$; Wilcoxon signed-rank for ChR2: 5Hz HC, $p = 0.0012207$; 5Hz LT, $p = 0.10986$; 20 Hz HC, $p = 0.01709$; 20 Hz LT, $p = 0.080322$; 50 Hz HC, $p = 0.01709$; 50Hz LT, $p = 0.00024414$.

**f,** Theta score across speed bins in ChR2 linear track sessions (5, 20, 50 Hz); shaded areas show mean ±SEM. 5 Hz, two-way within-subject RM ANOVA (stimulation x speed): $F(1,11) = 13.6340$, $p = 0.0035$; main effect of stimulation: $F(1,11) = 12.8109$, $p = 0.0043$. 20 Hz: stimulation x speed $F(1,11) = 2.4526$, $p = 0.1456$; stimulation $F(1,11) = 2.5372$, $p = 0.1395$. 50 Hz: stimulation x speed $F(1,11) = 0.0146$, $p = 0.9059$; stimulation $F(1,11) = 0.0125$, $p = 0.913$.

**g,** LFP spectral phase variance in non-theta and theta under urethane, showing that hippocampal LFP is phase locked to ChR2 stimulation frequency, marked by decreased variance.

**h,** Coupling of MS follower and pacemaker neurons to ChR2 stimulation frequency under urethane, quantified as the mean vector length of the preferred hippocampal phase distribution .Wilcoxon signed-rank tests (control vs stimulation): 5 Hz: non-theta, followers (F), $p = 0.0108437$; pacemakers (P), $p = 0.02148438$; theta, F, $p = 0.4631068$; P, $p = 0.1015625$; 20 Hz: non-theta, F, $p = 0.006713867$; P, $p = 0.0004882812$; theta, F, $p = 6.103516\times10^{-5}$; P, $p = 0.001953125$; 50 Hz: non-theta, F, $p = 0.8393555$; P, $p = 0.01367188$; theta, F, $p = 0.1530762$; P, $p = 0.0078125$.

**i,** LFP spectral phase variance in home cage and linear track, showing that hippocampal LFP is phase locked to ChR2 stimulation frequency.

**j,** Coupling of MS follower and pacemaker neurons to ChR2 stimulation frequency in free behavior, quantified as the mean vector length of the preferred hippocampal phase distribution. Wilcoxon signed-rank tests (control vs stimulation): 5 Hz: HC, F, $p = 2.1049\times10^{-7}$; P, $p = 0.00020675$; LT, F, $p = 7.8197\times10^{-8}$; P, $p = 4.0254\times10^{-5}$; 20 Hz: HC, F, $p = 3.0299\times10^{-8}$; P, $p = 2.7016\times10^{-5}$; LT, F, $p = 2.425\times10^{-8}$; P, $p = 2.7016\times10^{-5}$; 50 Hz: HC, F, $p = 2.425\times10^{-8}$; P, $p = 3.0884\times10^{-5}$; LT, F, $p = 2.8136\times10^{-8}$; P, $p = 2.7016\times10^{-5}$..

**k,** Polar histograms of theta phase preference of rhythmic MS neurons (followers and pacemakers) in linear track HS blockade, including only significantly modulated neurons ($p < 0.05$ in control periods). Sample sizes ($p < 0.05$): F, n = 46, P, n = 52.

**l-m,** Mean vector length of the preferred hippocampal theta phase distribution for all neurons (l, MS followers and pacemakers; m, hippocampal neurons; including both significantly and non-significantly modulated neurons) in HS blockade. Wilcoxon signed-rank tests: l, HC, F, p = 0.87727, P, p = 0.25385; LT, F, p = 0.35363, P, p = 0.00721; m, HC, p = 0.6536916; LT, p = 0.276053.

### Extended Data Figure 1.

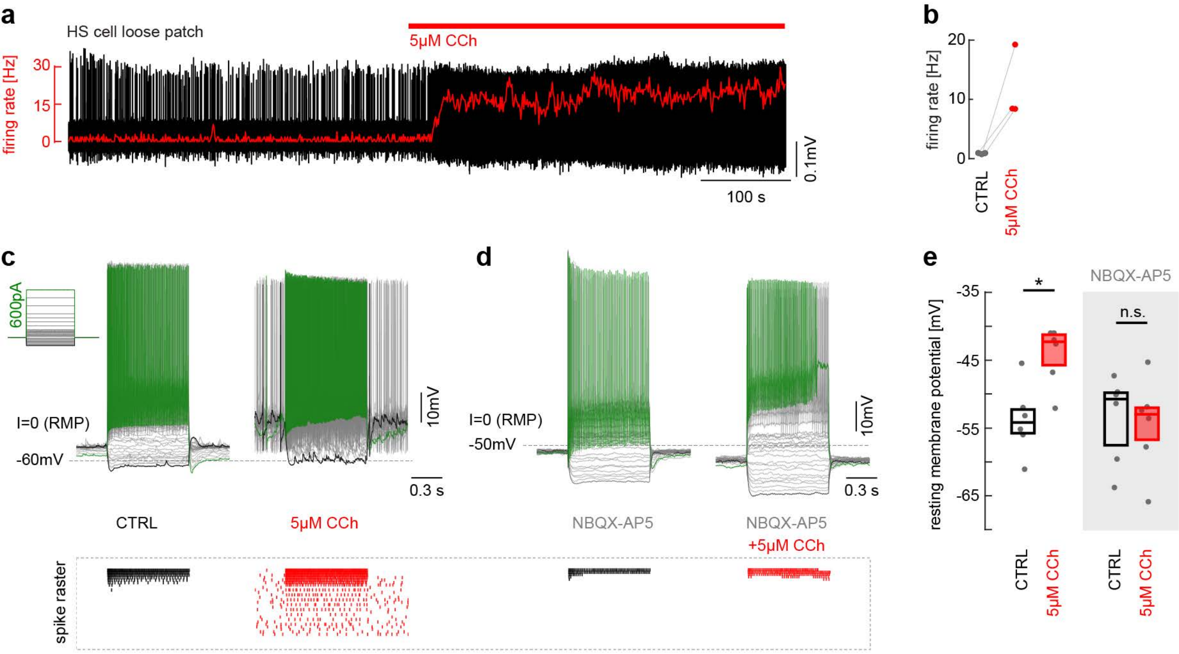

Extended Data Figure 2.

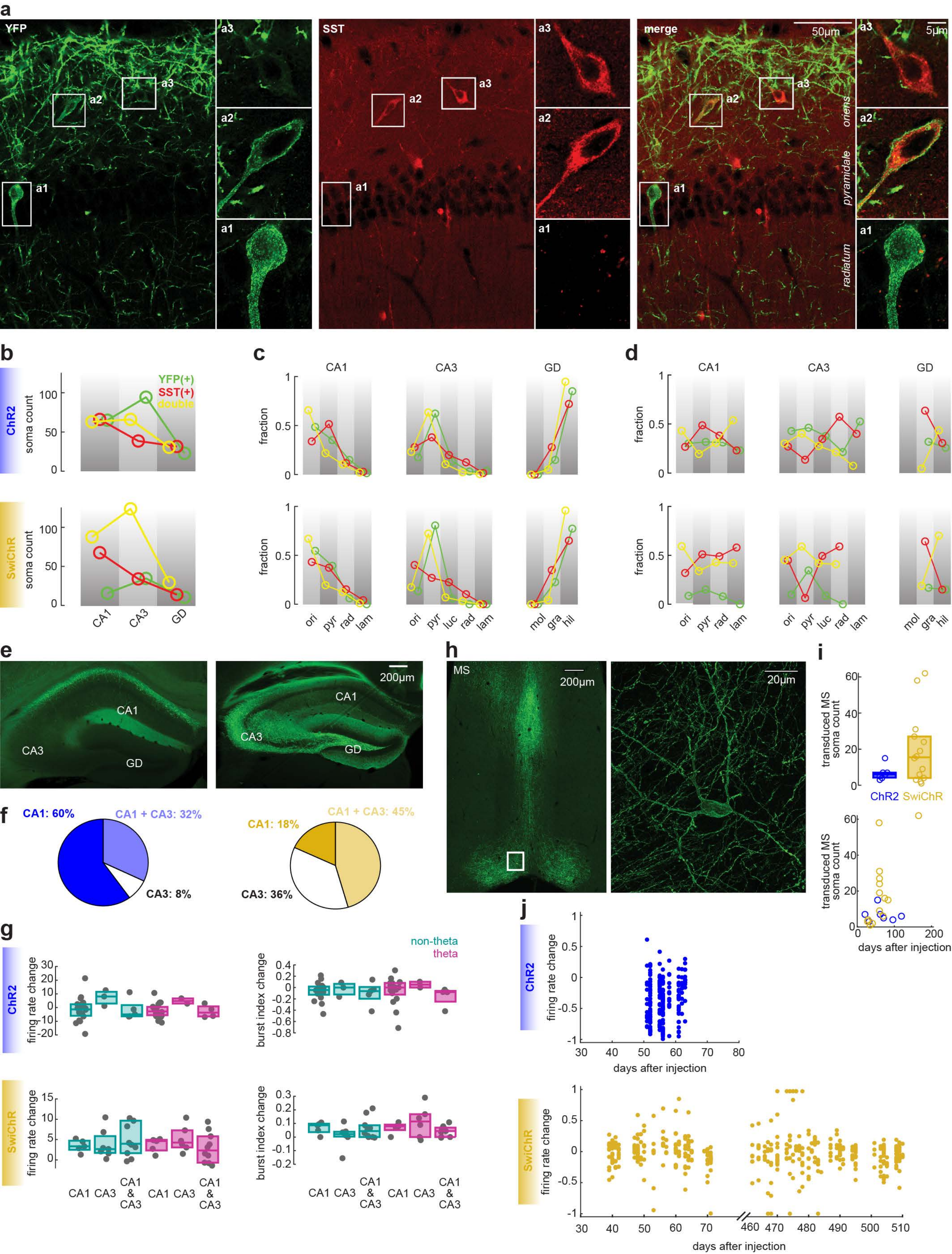

### Extended Data Figure 3.

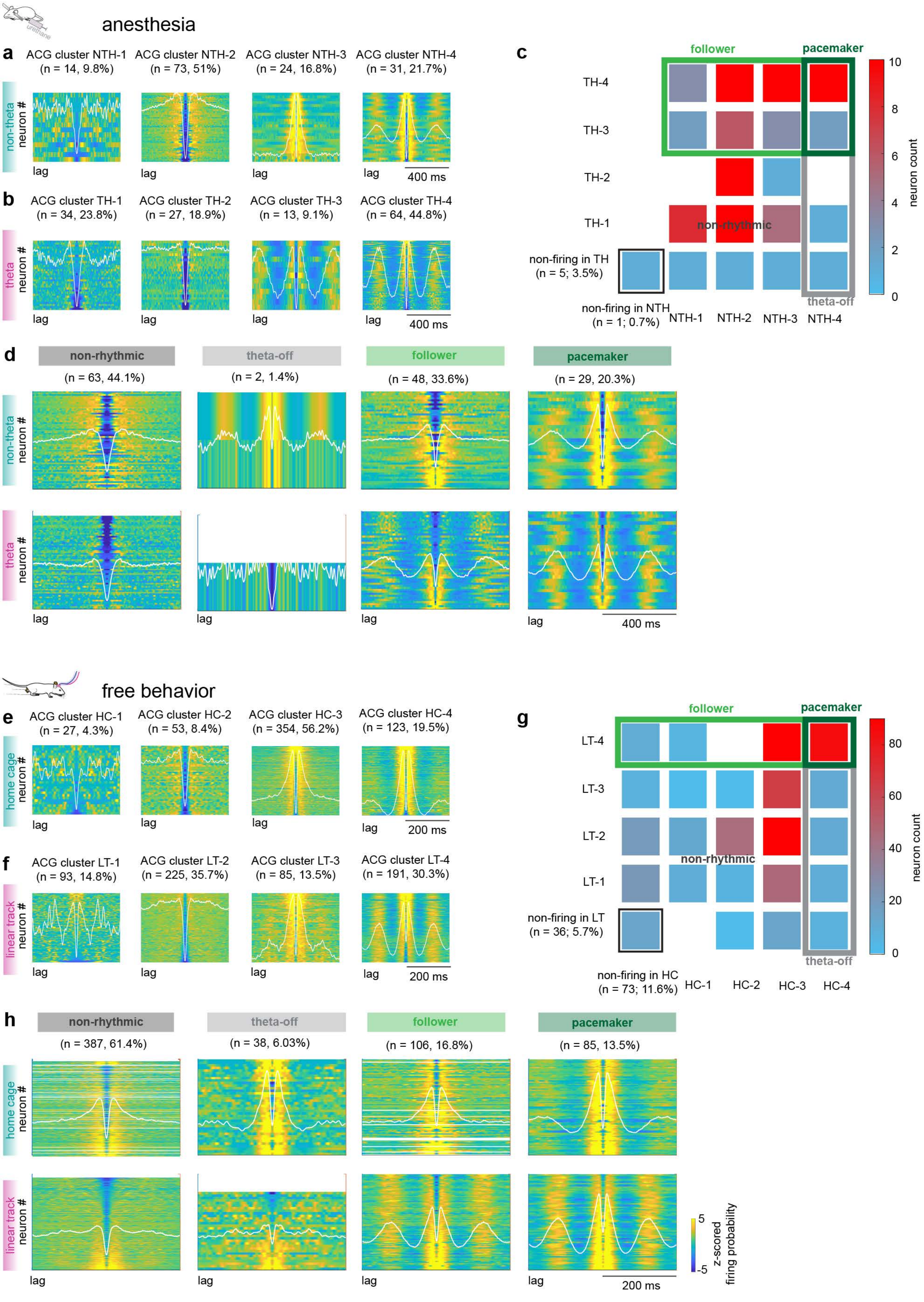

### Extended Data Figure 4.

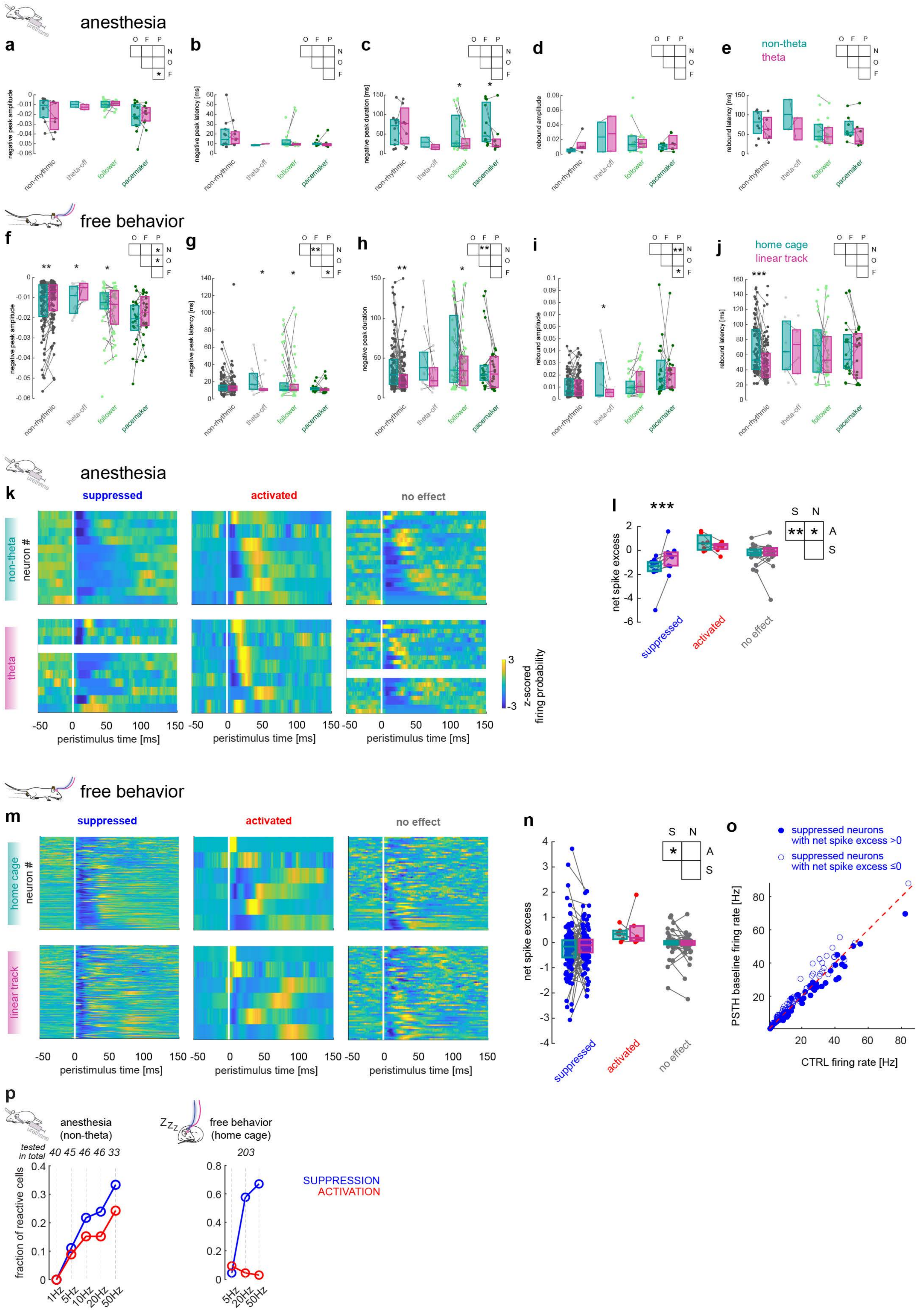

Extended Data Figure 5.

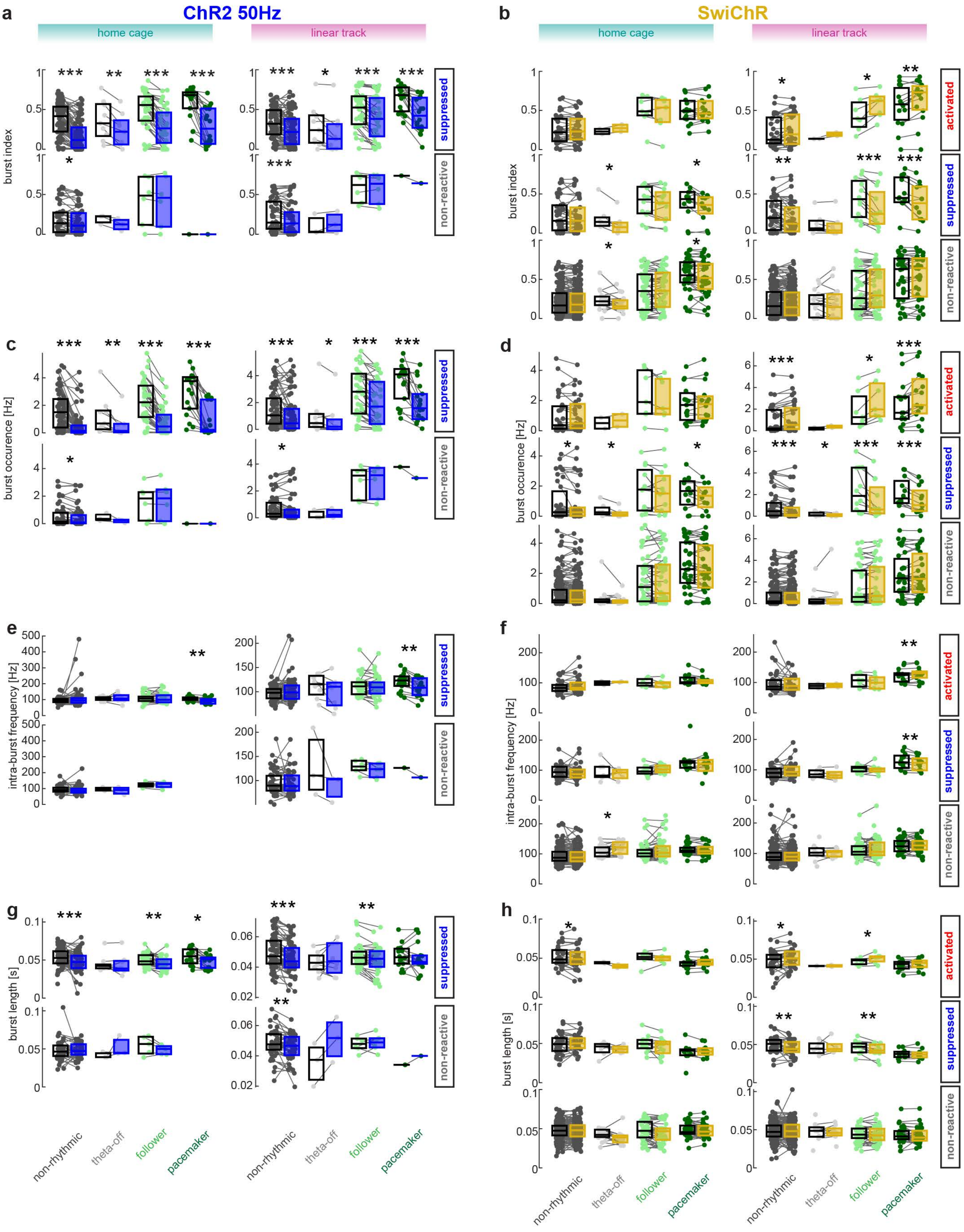

Extended Data Figure 6.

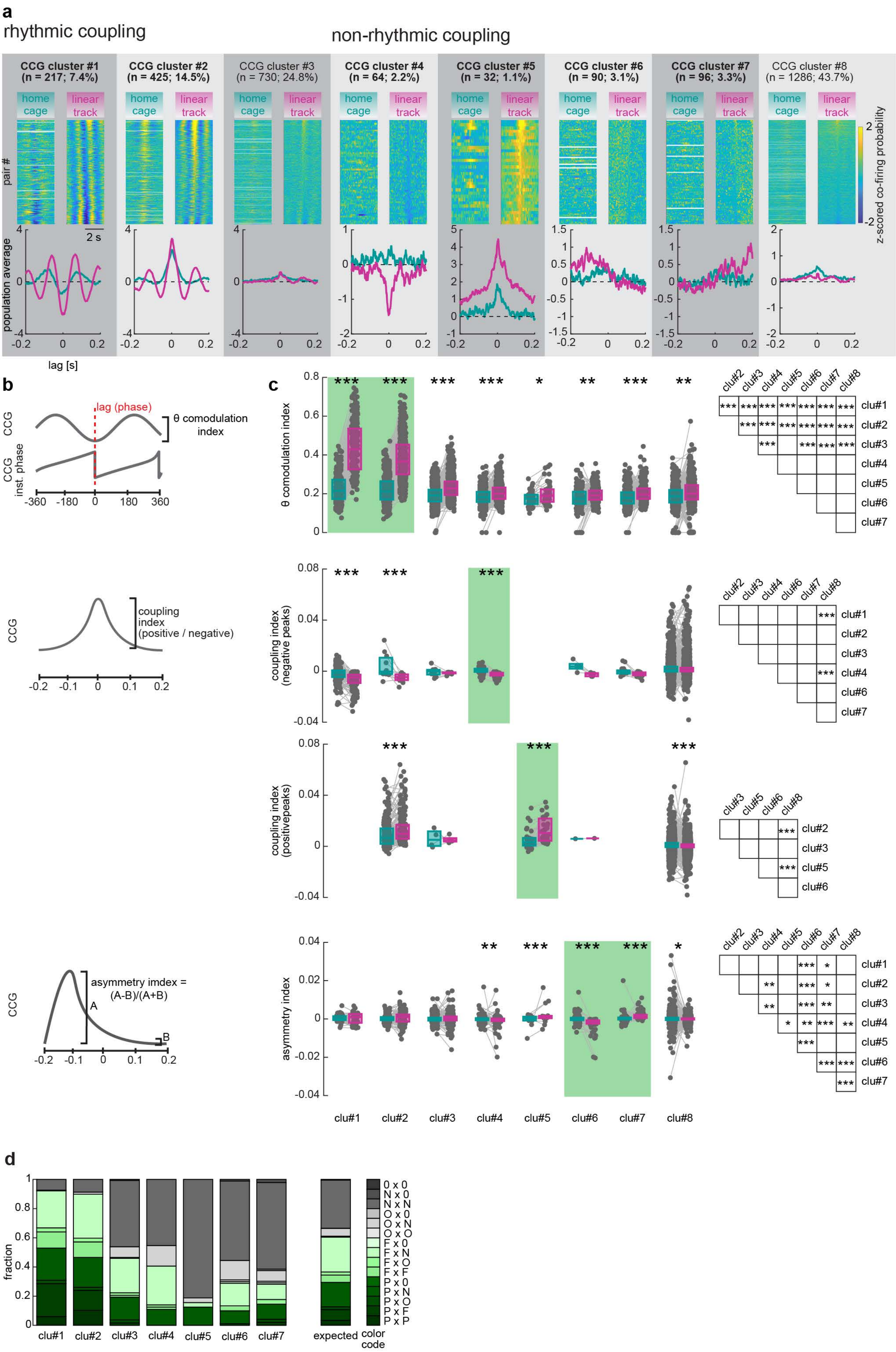

Extended Data Figure 7.

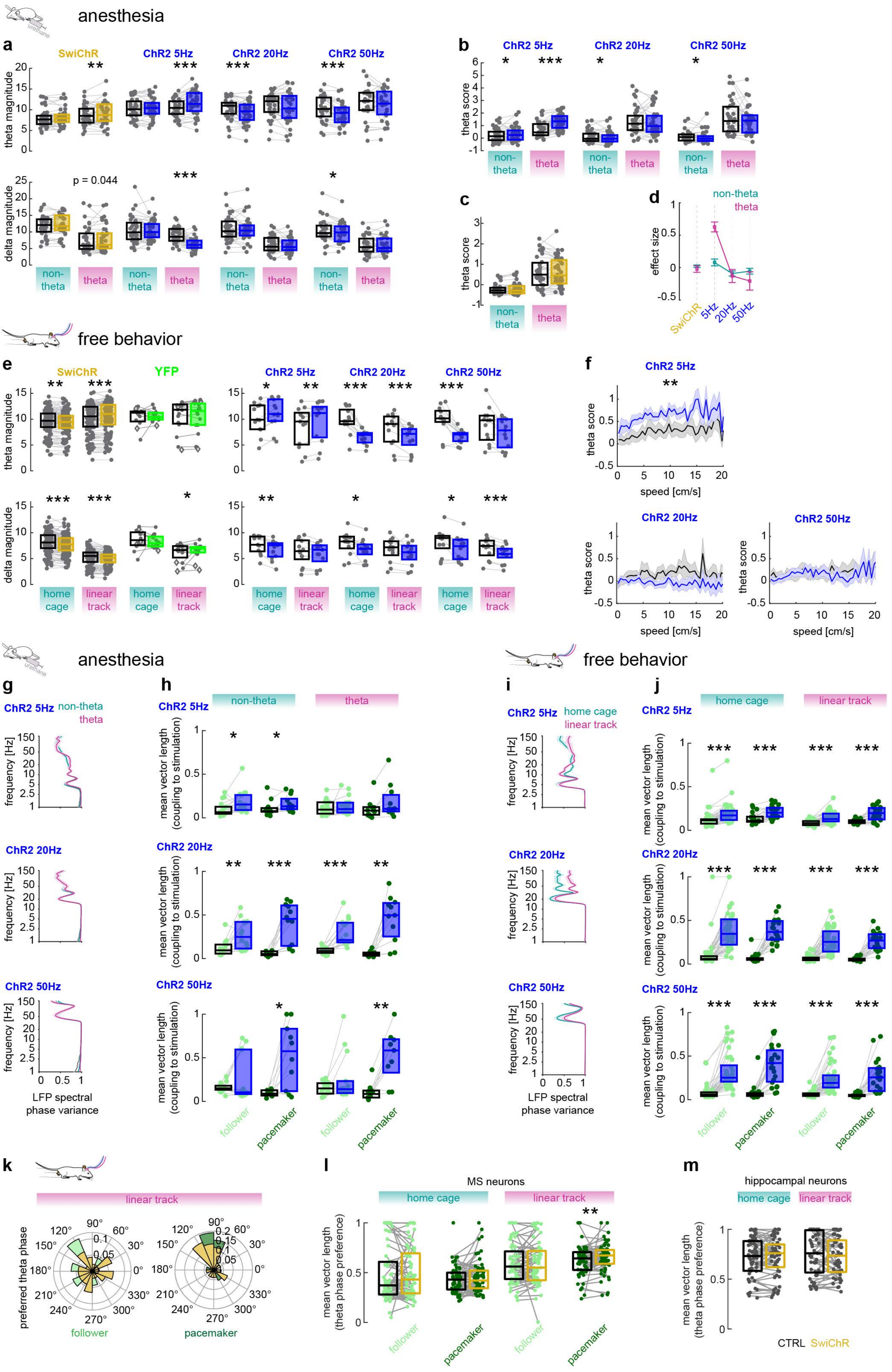

| | | rhythmicity category | | | |
|---|---|---|---|---|---|
| | **non-firing** | **non-rhythmic** | **theta-off** | **follower** | **pacemaker** |
| **ChR2 animals (n = 80)** | | | | | |
| recorded cells | 1 | 25 | 2 | 21 | 17 |
| labelled cells | 0 | 18 | 2 | 15 | 12 |
| recovered cells | 0 | 8 | 2 | 9 | 6 |
| clear PV immuno | 0 | 5 | 0 | 3 | 5 |
| PV positive | 0 | 0 | 0 | 1 | 5 |
| **SwiChR animals (n = 41)** | | | | | |
| recorded cells | 0 | 23 | 0 | 14 | 3 |
| labelled cells | 0 | 22 | 0 | 10 | 1 |
| recovered cells | 0 | 9 | 0 | 4 | 0 |
| clear PV immuno | 0 | 4 | 0 | 1 | 0 |
| PV positive | 0 | 0 | 0 | 0 | 0 |
| **Halo/Arch/eArch/ArchT animals (n = 35)** | | | | | |
| recorded cells | 0 | 15 | 0 | 13 | 9 |
| labelled cells | 0 | 8 | 0 | 9 | 5 |
| recovered cells | 0 | 2 | 0 | 3 | 1 |
| clear PV immuno | 0 | 1 | 0 | 2 | 1 |
| PV positive | 0 | 0 | 0 | 0 | 1 |
| **total PV immunopositivity** | | | | | |
| | **0** | **0/10** | **0** | **1/6** | **6/6** |

| animal ID | implanted probe into hippocampus | implanted probe into medial septum | analyzed sessions | analyzed units | non-firings (in the selected files) | non-rhythmics | theta-offs | followers | pacemakers | implanted optic fibers | notes |
|---|---|---|---|---|---|---|---|---|---|---|---|
| ***SwiChR animals for unit isolation*** | | | | | | | | | | | |
| 1814 | A1x32-10mm-100-177 | Buzsaki32L | 5 | 45 | 0 | 26 (57.8%) | 2 (4.4%) | 13 (28.9%) | 4 (8.9%) | 1 per each MS shank | no lfp analysis |
| 1833 | A1x32-10mm-100-177 | Buzsaki32L | 1 | 2 | 0 | 1 (50.0%) | 0 | 0 | 1 (50.0%) | 1 per each MS shank | no lfp analysis |
| 1865 | A1x32-10mm-100-177 | Buzsaki32L | 15 | 100 | 0 | 76 (76.0%) | 2 (2.0%) | 11 (11.0%) | 11 (11.0%) | 1 per each MS shank | no lfp analysis |
| 1869 | Buzsaki32L - only 1 shank | Buzsaki32L | 3 | 21 | 0 | 14 (66.7%) | 0 | 2 (9.5%) | 5 (23.8%) | 1 per each MS shank | |
| 1966 | A1x32-10mm-100-177 | Buzsaki32L | 3 | 19 | 1 (5.3%) | 17 (89.5%) | 1 (5.3%) | 0 | 0 | 1 per each MS shank | ripple analysis performed |
| 1988 | A1x32-10mm-100-177 | Buzsaki32L | 29 | 169 | 13 (7.7%) | 78 (46.2%) | 20 (11.8%) | 24 (14.2%) | 34 (20.1%) | 1 per each MS shank & blind | ripple analysis performed |
| 1989 | Buzsaki32L - only 3 shanks | Buzsaki32L | 14 | 71 | 0 | 48 (67.6%) | 1 (1.4%) | 15 (21.1%) | 7 (9.9%) | 1 per each MS shank | |
| ***SwiChR animals for ripple analysis*** | | | | | | | | | | | |
| 1965 | A4x16-6mm-50-200-177 & A1x32-10mm-100-177 | | 1 | | | | | | | bilaterally into MS | ripple analysis performed |
| 1982 | A4x16-6mm-50-200-177 & A1x32-10mm-100-177 | | 1 | | | | | | | bilaterally into MS | ripple analysis performed |
| 1986 | A4x16-6mm-50-200-177 & A1x32-10mm-100-177 | | 1 | | | | | | | bilaterally into MS & blind | ripple analysis performed |
| 1993 | A4x16-6mm-50-200-177 & A1x32-10mm-100-177 | | 1 | | | | | | | bilaterally into MS & blind | ripple analysis performed |
| additional n = 7 animals were not used (died during or after surgery, broken silicon probe, bad quality units or probe not reaching MS) | | | | | | | | | | | |
| ***ChR2 animals for unit isolation*** | | | | | | | | | | | |
| 1695 | A1x32-10mm-100-177 | Buzsaki32L | 7 | 112 | 0 | 68 (60.7%) | 5 (4.5%) | 23 (20.5%) | 16 (14.3%) | fimbria | |
| 1750 | A1x32-10mm-100-177 | Buzsaki32L | 6 | 91 | 0 | 59 (64.8%) | 7 (7.7%) | 18 (19.8%) | 7 (7.7%) | fimbria | |
| additional n = 8 animals were not used (died during or after surgery, broken silicon probe, bad quality units or probe not reaching MS) | | | | | | | | | | | |
| ***YFP animals for ripple and LFP analysis*** | | | | | | | | | | | |
| 1946 | A1x32-10mm-100-177 | | 6(LT), 2(HC) | | | | | | | bilaterally into MS | ripple analysis performed |
| 1969 | A1x32-10mm-100-177 | | 3 | | | | | | | bilaterally into MS | ripple analysis performed |
| 1972 | A4x16-6mm-50-200-177 & A1x32-10mm-100-177 | | 2 | | | | | | | bilaterally into MS | ripple analysis performed |
| 1977 | A1x32-10mm-100-177 | | 2 | | | | | | | bilaterally into MS | ripple analysis performed |
| additional n = 3 animals were not used (died after surgery, bad injection or broken silicon probe) | | | | | | | | | | | |
